# Detection of a historic reservoir of bedaquiline / clofazimine resistance associated variants in *Mycobacterium tuberculosis*

**DOI:** 10.1101/2020.10.06.328799

**Authors:** Camus Nimmo, Arturo Torres Ortiz, Juanita Pang, Mislav Acman, Cedric C.S. Tan, James Millard, Nesri Padayatchi, Alison Grant, Max O’Donnell, Alex Pym, Ola B Brynildsrud, Vegard Eldholm, Louis Grandjean, Xavier Didelot, François Balloux, Lucy van Dorp

## Abstract

Drug resistance in tuberculosis (TB) poses a major ongoing challenge to public health. The recent inclusion of bedaquiline into TB drug regimens has improved treatment outcomes, but this advance is threatened by the emergence of strains of *Mycobacterium tuberculosis* (*Mtb*) resistant to bedaquiline. Clinical bedaquiline resistance is most frequently conferred by off-target resistance-associated variants (RAVs) in the *mmpR5* gene (*Rv0678*), the regulator of an efflux pump, which can also confer cross-resistance to clofazimine, another TB drug. We compiled a dataset of 3,682 *Mtb* genomes, including 150 carrying variants in *mmpR5* that have been associated to borderline (henceforth intermediate) or confirmed resistance to bedaquiline. We identified eight cases where RAVs were present in the genomes of strains collected prior to the use of bedaquiline in TB treatment regimes. Phylogenetic reconstruction points to multiple emergence events and circulation of RAVs in *mmpR5*, some estimated to predate the introduction of bedaquiline. However, epistatic interactions can complicate bedaquiline drug-susceptibility prediction from genetic sequence data. Indeed, in one clade of isolates where the RAV Ile67fs is estimated to have emerged prior to the antibiotic era, co-occurrence of mutations in *mmpL5* are found to neutralise bedaquiline resistance. The presence of a pre-existing reservoir of *Mtb* strains carrying bedaquiline RAVs prior to its clinical use augments the need for rapid drug susceptibility testing and individualised regimen selection to safeguard the use of bedaquiline in TB care and control.

## Introduction

Drug-resistant tuberculosis (DR-TB) currently accounts for 450,000 of the 10 million new tuberculosis (TB) cases reported annually^1^. Treatment outcomes for multidrug-resistant TB (MDR-TB), resistant to at least rifampicin and isoniazid, have historically been poor, with treatment success rates of only 50-60% in routine programmatic settings^2,3^. The discovery of bedaquiline, a diarylquinoline antimycobacterial active against ATP synthase, which is highly effective against *Mycobacterium tuberculosis* (*Mtb*)^4^, was reported in 2004. Following clinical trials, which confirmed reduced time to culture conversion in patients with DR-TB^5^, in 2012 bedaquiline received an accelerated Food and Drug Administration (FDA) licence for use in DR-TB^6^.

Cohort studies of patients treated with bedaquiline-containing regimens against MDR-TB report success rates of 70-80%^7,8^. Similar results have been achieved for extensively drug-resistant TB (XDR-TB, traditionally defined as MDR-TB strains with additional resistance to fluoroquinolones and injectables), where treatment outcomes without bedaquiline are even worse^9,10^. In light of these promising results, the World Health Organization (WHO) now recommends that bedaquiline be included in all MDR-TB regimens^11^. It has played a central role in the highly successful ZeNix^12^ and TB-PRACTECAL^13^ trials of bedaquiline, pretomanid and linezolid (+/− moxifloxacin) six-month all-oral regimens for DR-TB. These are now incorporated in WHO guidance. In addition, bedaquiline is positioned as a key drug in multiple phase III clinical trials for drug-susceptible TB (SimpliciTB, ClinicalTrials.gov NCT03338621; TRUNCATE-TB^14^).

Resistance in *Mtb* is typically reported shortly after the introduction of a novel TB drug and often appears sequentially^15,16^. For example, mutations conferring resistance to isoniazid – one of the first antimycobacterials – tend to emerge prior to resistance to rifampicin, the other major first-line drug. These also predate resistance mutations to second-line drugs, so termed because they are used clinically to treat patients infected with strains already resistant to first-line drugs. This was observed, for example, in KwaZulu-Natal, South Africa, where resistance-associated mutations accumulated over decades prior to their identification, leading to a major outbreak of extensively drug-resistant TB (XDR-TB)^16^. Unlike other major drug-resistant bacteria, *Mtb* reproduces strictly clonally and systematically acquires resistance by chromosomal mutations rather than via horizontal gene transfer or recombination^17^. This allows phylogenetic reconstructions, based on whole genome sequencing data, to be used to infer the timings of emergence and subsequent spread of variants in *Mtb* that have been suggested to reduce drug susceptibility, termed resistance-associated variants (RAVs).

A number of mechanisms have been implicated in conferring bedaquiline resistance. For example, mutations conferring resistance have been selected *in vitro*, located in the *atpE* gene encoding the F1F0 ATP synthase, the target of bedaquiline^18^. Off-target resistance-conferring mutations have also been found in *pepQ* in a murine model and potentially in a small number of patients^19^. However, the primary mechanism of resistance observed in clinical isolates has been identified in the context of off-target resistance-associated variants (RAVs) in the *mmpR5 (Rv0678)* gene, a negative repressor of expression of the MmpL5 efflux pump. Loss of function of mmpR5 leads to pump overexpression^20^ and increased minimum inhibitory concentrations (MIC) to bedaquiline, along with the recently repurposed antimycobacterial clofazimine, fusidic acid, the azole class of antifungal drugs (which also have antimycobacterial activity), as well as to the novel therapeutic class of DprE1 inhibitors in clinical trials^21,22^. Aligned with this mechanism of resistance, coincident mutations leading to loss of function of the MmpL5 efflux pump can negate the resistance-inducing effect of MmpR5 loss of function^23^.

A range of single nucleotide polymorphisms (SNPs) and frameshift *mmpR5* mutations have been associated with resistance to bedaquiline and are often present as heteroresistant alleles in patients^24–34^. In contrast to most other RAVs in *Mtb,* which often cause many-fold increases in MIC and clear-cut resistance, *mmpR5* variants may be associated with normal MICs or subtle increases in bedaquiline MIC, although they may still be clinically important^35^. These increases may not cross the current WHO critical concentrations used to classify resistant versus susceptible strains (0.25µg/mL on Middlebrook 7H11 agar, or 1µg/mL in Mycobacteria Growth Indicator Tube [MGIT] liquid media). The first version of the WHO tuberculosis drug resistance catalogue does not contain any bedaquiline RAVs, although a subsequent meta-analysis identified two RAVs (a*tpE* Ala63Pro and *mmpR5* Ile67fs)^34^. Bedaquiline has a long terminal half-life of up to 5.5 months^6^, leading to the possibility of subtherapeutic concentrations where adherence is suboptimal or treatment is interrupted, which could act as a further driver of resistance.

Bedaquiline and clofazimine cross-resistance has now been reported across three continents following the rapid expansion in usage of both drugs^25,30,36,37^, and is associated in some cases with poor adherence to therapy and inadequate regimens. However, baseline isolates in 8/347 (2.3%) patients from phase IIb bedaquiline trials demonstrated *mmpR5* RAVs and high bedaquiline MICs in the absence of prior documented use of bedaquiline or clofazimine^38^. This suggests that bedaquiline RAVs may have been in circulation prior to the usage of either of these drugs, which may be expected in the case of mutations which do not have major fitness consequences^39^. While there have been isolated clinical reports from multiple geographical regions, the global extent of bedaquiline resistance emergence and spread has not yet been investigated.

In this study, we characterise and date the emergence of variants in *mmpR5*, including those implicated as bedaquiline RAVs, in the two global *Mtb* lineage 2 (L2) and lineage 4 (L4) lineages, which include the majority of drug resistance strains^15^. Phylogenetic analyses of two datasets comprising 1,514 *Mtb* L2 and 2,168 L4 whole genome sequences revealed the emergence and spread of multiple *mmpR5* variants associated to resistance or borderline (intermediate) resistance to bedaquiline prior to its first clinical use. This pre-existing reservoir of bedaquiline/clofazimine-resistant *Mtb* strains suggests *mmpR5* RAVs exert a relatively low fitness cost which could be rapidly selected for as bedaquiline and clofazimine are more widely used in the treatment of TB.

## Results

### The global diversity of *Mtb* lineage L2 and L4

To investigate the global distribution of *Mtb* isolates with variants in *mmpR5*, we curated two large datasets of whole genomes from the two dominant global lineages L2 and L4. Both datasets were enriched for samples with variants in *mmpR5* following a screen for variants in public sequencing repositories (see **Methods**) and retaining those samples uploaded with accompanying full metadata for geolocation and time of sampling (**Figure 1, Supplementary Table S1-S2, Supplementary Figure S1)**. The final L2 dataset included 1,514 isolates collected over 24.5 years (between 1994 and 2019) yielding 29,205 SNPs. The final L4 dataset comprised 2,168 sequences collected over 232 years, including three samples from 18^th^ century Hungarian mummies^40^, encompassing 67,585 SNPs. Both datasets included recently generated data from South Africa (155 L2, 243 L4)^41,42^ and new whole genome sequencing data from Peru (9 L2, 154 L4).

**Figure 1:**
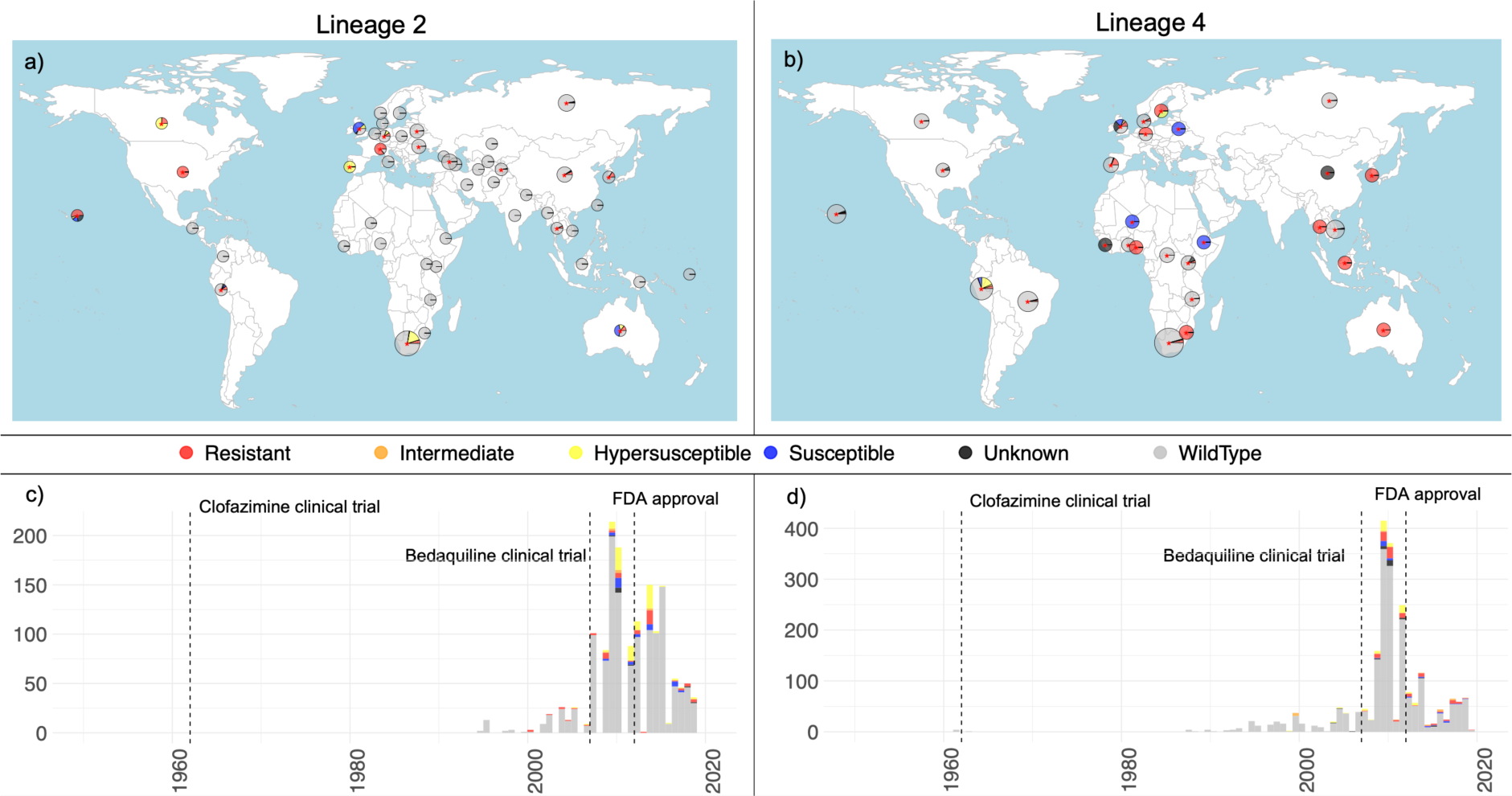
Compiled global *Mtb* genomic datasets. Panels a) and b) provide the geographic location of isolates included in the lineage 2 and lineage 4 datasets respectively. Pies are scaled by the number of samples per country (raw data available in **Supplementary Table S1**) with the colours providing the fraction of genomes with any nonsynonymous/frameshift variants detected in *mmpR5* (coloured as per the legend). Countries comprising samples with known RAVs are highlighted with a red asterisk. Genomic data for which no associated metadata on the geographic location of sampling was available are shown in the Pacific Ocean. Panels c) and d) provide the collection dates associated to each genome in the lineage 2 and lineage 4 datasets respectively highlighting those with any variants in *mmpR5* (colour, as per legend). Lineage 4 *Mtb* obtained from 18^th^ century mummies are excluded from this plot but included in all analyses. The vertical dashed lines indicate the dates of the first clinical trials for clofazimine, bedaquiline and FDA approval of bedaquiline for clinical use.

Consistent with previous studies^43–45^, both datasets are highly diverse and exhibit strong geographic structure (**Figure 2**). As a nonrecombining clonal organism, identification of mutations in *Mtb* can provide a mechanism to predict phenotypic resistance from a known panel of genotypes^46,47^. Based on genotypic profiling^47^, 911 strains within the L2 dataset were classified as MDR-TB (60%) and 295 (20%) as XDR-TB. Within the L4 dataset, 911 isolates were classified as MDR-TB (42%) and 115 as XDR-TB (5%). The full phylogenetic distribution of resistance profiles is provided in **Supplementary Figure S2**. As is commonplace with genomic datasets, the proportion of drug-resistant strains exceeds their actual prevalence, due to the overrepresentation of drug-resistant isolates in public sequencing repositories.

**Figure 2:**
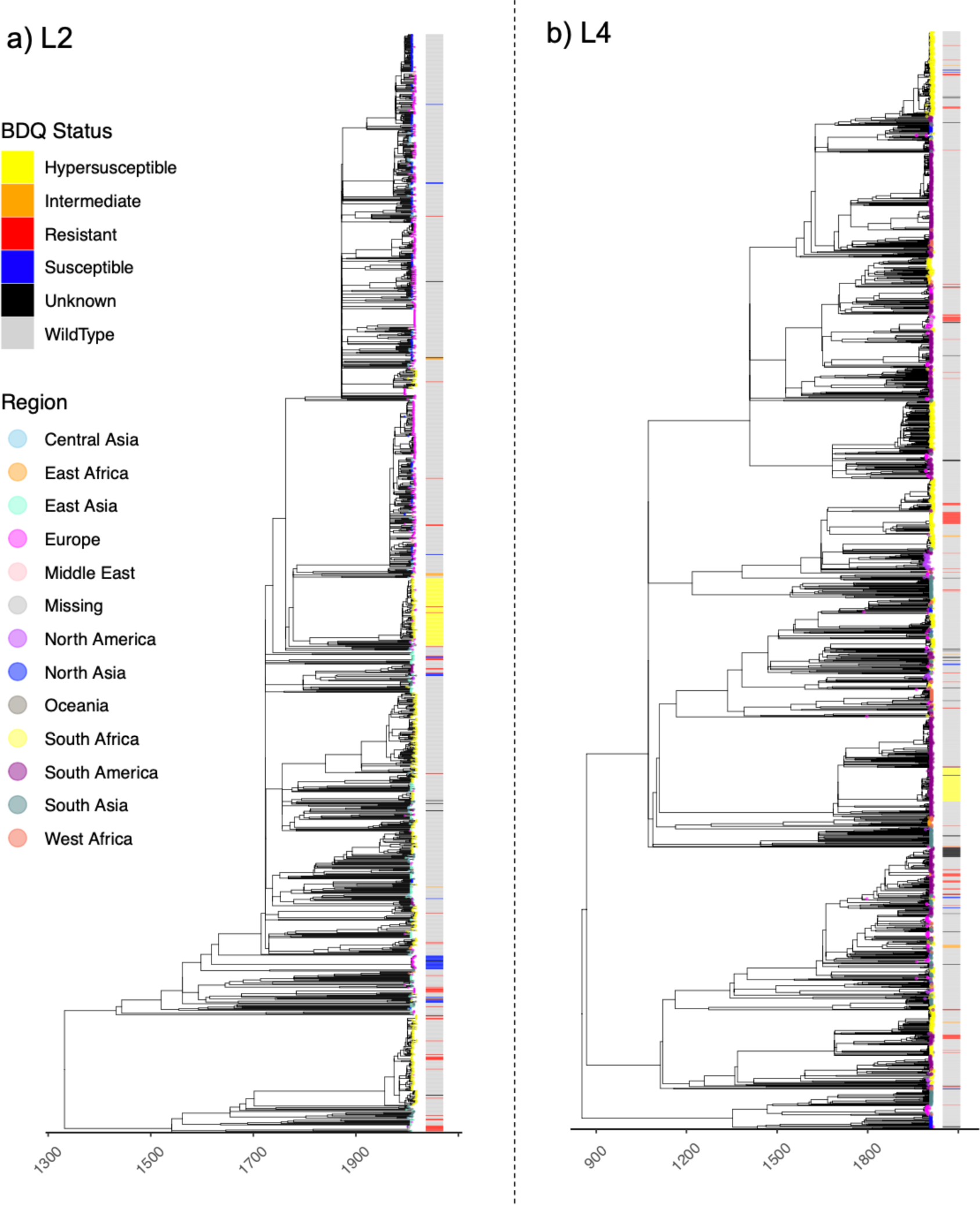
Global time calibrated *Mtb* phylogenies. Inferred dated phylogenies (x-axis) for the a) lineage 2 and b) lineage 4 datasets. Tips are coloured by the geographic region of sampling as given in the legend. The bar provides the assessed phenotype (colour) based on assignment of nonsynonymous/frameshift variants in *mmpR5*.

Both the L2 and L4 phylogenetic trees displayed a significant temporal signal following date randomisation (**Supplementary Figure S3**), making them suitable for time-calibrated phylogenetic inference^48^. We estimated the time to the Most Recent Common Ancestor (tMRCA) of both datasets using a Bayesian tip-dating analysis (BEAST2) run on a representative subset of genomes from each dataset (see **Methods**, **Supplementary Table 3**, **Supplementary Figure S4**). For the final temporal calibration of the L2 dataset we applied an estimated clock rate of 7.7×10^−8^ (4.9×10^−8^ - 1.03×10^−7^) substitutions per site per year, obtained from the subsampled BEAST2^48^ analysis, to the global maximum likelihood phylogenetic tree. This resulted in an estimated tMRCA of 1332CE (945CE-1503CE). Using the same approach for the L4 dataset we estimated a clock rate of 7.1×10^−8^ (6.2×10^−8^ - 7.9×10^−8^) substitutions per site per year resulting in an estimated tMRCA of 853CE (685CE – 967CE) (**Figure 2**). We observed a slightly higher, yet statistically not significant, clock rate in L2 compared to L4 (**Supplementary Table S3**), with all estimated substitution rates falling largely in line with previously published estimates^49^.

### Identification of variants in *mmpR5*

Since *atpE* and *pepQ* bedaquiline RAVs are found at low prevalence (1 L2 isolate [0.03%] and 18 L4 isolates [0.49%]), we focused on characterising mutations in *mmpR5*. In total we identified the presence of non-synonymous and promoter *mmpR5* variants in 437 sequences (193 L2 [12.8%], 244 L4 [11.3%]). We classified all identified non-synonymous and promoter mutations in *mmpR5*, based on an evaluation of their phenotypic impact through review of published literature, into six phenotypic categories for bedaquiline susceptibility: wild type, hypersusceptible, susceptible, intermediate, resistant, and unknown (full references available in **Supplementary Table S4, Supplementary Figures S5-S7)**. Across both lineages, 148 sequences were considered as bedaquiline resistant (i.e., classified as intermediate or resistant). The most frequently observed variants are listed in **Table 1**.

**Table 1:** Frequency of all *mmpR5* variants occurring ≥ 5 times in dataset and associated resistance classification. *Where co-existing *mmpL5* mutations were identified this is indicated – only one mutation was found (*mmpL5* Arg202fs) and it was in the presence of *mmpR5* Ile67fs mutations only, with no other co-existing *mmpR5* variants.

| Variant | Associated phenotype | L2 | L4 | Total |
| --- | --- | --- | --- | --- |
| C-11A | Hypersusceptible | 93 |  | 93 |
| Ile67fs + mmpL5 Arg202fs | Hypersusceptible |  | 66 | 66 |
| Asp5Gly | Susceptible | 20 | 3 | 23 |
| Met146Thr | Resistant | 2 | 20 | 22 |
| Ile67fs | Resistant | 5 | 17 | 22 |
| Leu40Val | Susceptible |  | 19 | 19 |
| Arg90Cys | Intermediate | 2 | 9 | 11 |
| Glu49fs | Resistant | 2 | 8 | 10 |
| Val20Ala | Intermediate | 1 | 6 | 7 |
| Ala59Val | Resistant | 7 |  | 7 |
| Val1Ala | Resistant | 6 |  | 6 |
| Gly121Arg | Resistant | 5 |  | 6 |
| Asp141fs | Unknown |  | 6 | 6 |
| Asn98Asp | Resistant |  | 6 | 6 |
| Arg96Gly | Resistant |  | 5 | 5 |
| Arg109Leu + Arg156fs | Resistant |  | 5 | 5 |

We identified a significant relationship between the presence of *mmpR5* variants and drug resistance status in both the L2 and L4 datasets (**Supplementary Figure S8-S9**), though in both cases we identified otherwise fully phenotypically susceptible isolates carrying *mmpR5* RAVs. Notably we identified 24 sequenced isolates carrying nonsynonymous/frameshift variants in *mmpR5* uploaded with collection dates prior to the first clinical trials for bedaquiline in 2007. This comprised ten L2 isolates collected before 2007, of which eight harboured variants previously associated to phenotypic bedaquiline resistance (RAVs). For L4 we identified 15 sequences with *mmpR5* variants predating 2007, of which six have been previously classified as carrying mutations conferring a bedaquiline resistance phenotype above wild-type (’intermediate’) (**Figure 1c-d, Supplementary Table S5**).

Within the datasets, we identified one L2 isolate (ERR2677436 sampled in Germany in 2016) which already had two *mmpR5* RAVs at low allele frequency – Val7fs (11%) and Val20Phe (20%) – and also contained two low frequency *atpE* RAVs: Glu61Asp (3.2%) and Alal63Pro (3.7%)^50^. We also identified three isolates obtained in 2007-08 from separate but neighbouring Chinese provinces carrying the *Rv1979c* Val52Gly RAV, which has been suggested to be associated with clofazimine resistance in a study from China^25^ but was associated with a normal MIC in another^39^, with its role in resistance remaining unclear^31^. Furthermore, frameshift and premature stop mutations in *pepQ* have been previously associated with bedaquiline and clofazimine resistance. In this dataset, we identified 18 frameshift mutations in *pepQ* across 11 patients, one of which also had a *mmpR5* frameshift mutation. In one isolate the *pepQ* frameshift occurred at the Arg271 position previously reported to be associated with bedaquiline resistance^19^.

Sixty-three genomes harboured nonsynonymous *mmpR5* variants of unknown phenotypic effect (12 L2, 28 L4), corresponding to 23 unique mutations or combinations of mutations. To assess properties associated to RAVs which may be useful predictors of the phenotypic effect of these unknown variants we employed a machine learning approach, providing a foundation for further exploration of genomic features associated to RAV status (see **Supplementary Note 1**).

### The time to emergence of *mmpR5* variants

To estimate the age of the emergence of different *mmpR5* non-synonymous variants, we identified all nodes in each of the L2 and L4 global time calibrated phylogenies delineating clades of isolates carrying a particular *mmpR5* variant (**Figure 3, Supplementary Table S6, Supplementary Table S7**). For the L2 dataset we identified 58 unique phylogenetic nodes where *mmpR5* RAVs emerged, of which 40 were represented by a single genome. The point estimates for these nodes ranged from March 1845 to November 2018. Eight nonsynonymous/frameshift variants in *mmpR5*, including four bedaquiline RAVs (Met139Ile, Cys46fs, Ala59Val, Asn98fs) and one case expected to lead to an intermediate phenotype (Arg90Cys), were estimated to have emergence dates (point estimates) predating the first bedaquiline clinical trial in 2007 (**Supplementary Figure S10**).

**Figure 3:**
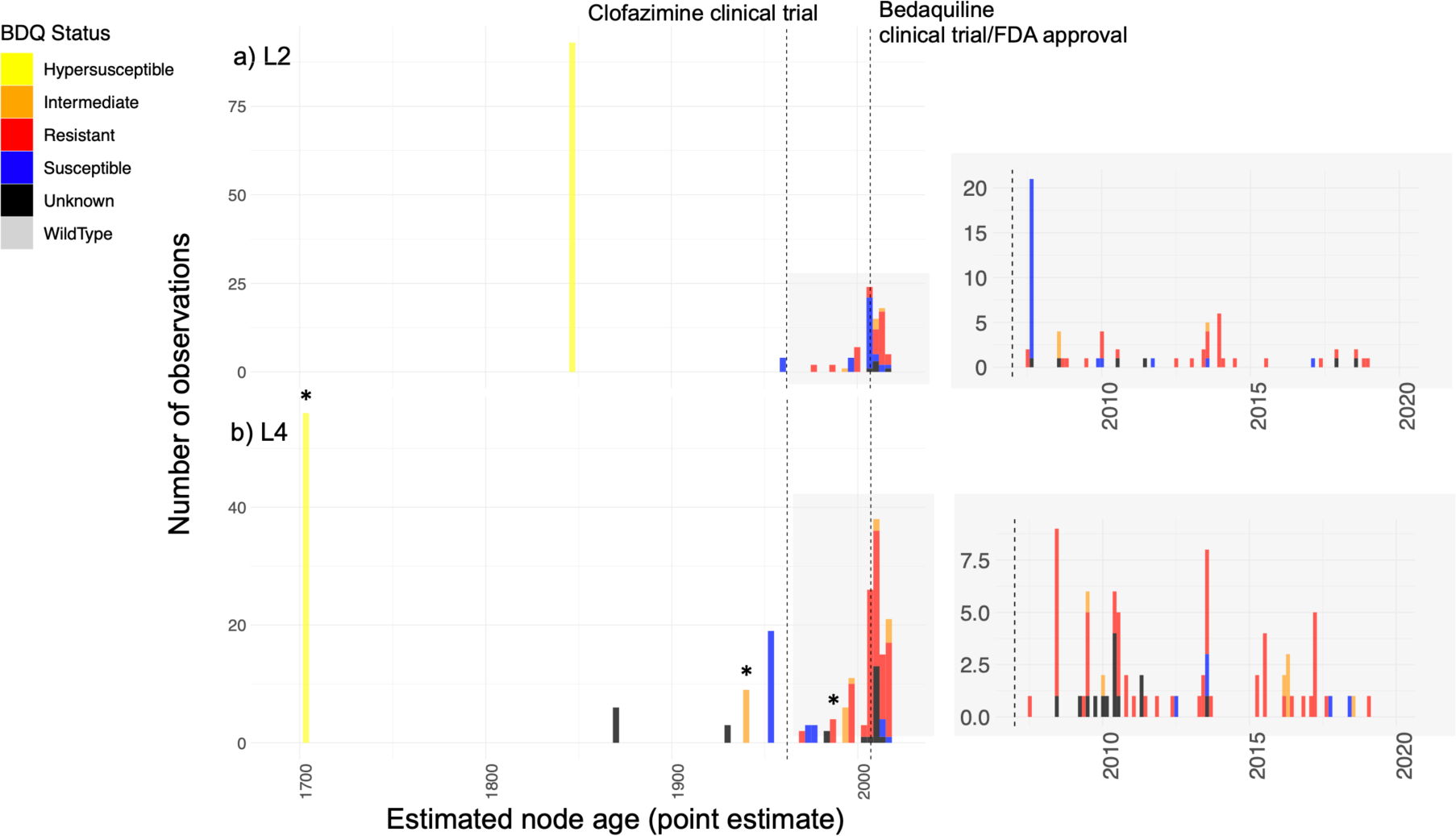
Estimated age of emergence of *mmpR5* nonsynonymous/frameshift variants. Inferred point estimates for the age of emergence of clades with *mmpR5* variants for the lineage 2 (a) and lineage 4 (b) datasets, including a zoomed in reproduction of the period from 2007-2020. Y-axis provides the absolute number of sequences descending from the identified and dated nodes. The *mmpR5* RAV status is given by the colour as defined in the legend at bottom. *indicates phenotypic data available for considered isolates that are supportive of MIC classification (see text).The full mutation timelines are provided in **Supplementary Figures 11-12** and **Supplementary Table S7**.

For the L4 dataset we identified 85 unique nodes where *mmpR5* RAVs emerged, of which 59 were represented by a single isolate in the dataset. The point estimates for these nodes ranged from September 1701 to January 2019 (**Figure 3, Supplementary Figure S11**). Nineteen *mmpR5* mutations, including six unique bedaquiline RAVs (Gln22Arg, Asn98Asp, Ile67fs x2, Arg96Gly, Met146Thr, Asn98Asp) and two predicted to have an intermediate phenotype (Arg90Cys, Ser53Leu), were estimated to have emerged prior to 2007. Arg90Cys, in particular was estimated to emerge between 1930-1947, suggesting the likely circulation of variants which lead to a response to bedaquiline above wild-type pre-existed the first clinical trials for clofazimine in the 1960s. While we identified no nodes with a second emergence of *mmpR5* nonsynonymous/frameshift mutations across the L4 dataset, eight nodes were identified in the L2 dataset where a clade already carrying a nonsynonymous/frameshift variant in *mmpR5* subsequently acquired a second nonsynonymous/frameshift mutation.

In the L4 dataset, we noted one large clade of 66 samples, predominantly collected in Peru (henceforth Peruvian clade), which all carry the Ile67fs *mmpR5* resistance associated mutation^36,50,51^. While it is not inconceivable that multiple independent emergences of Ile67fs occurred in this clade, the more parsimonious scenario is a single ancestral emergence. We estimate the time of this emergence to 1702 (1657-1732) (**Figure 3, Supplementary Figure S11-S12**). Of significance, we identified a frameshift mutation in the adjacent *MmpL5* efflux pump (Arg202fs) in isolates from this Peruvian clade, the protein whose overexpression mediates bedaquiline resistance following loss-of-function of the MmpR5 regulatory protein. This frameshift, which leads to a premature stop codon at amino acid 206, is expected to counteract the otherwise resistance-conferring mutation. This epistatic interaction restoring bedaquiline susceptibility has recently been described elsewhere^23,39^. The *mmpL5* frameshift mutation was present in all isolates in the Peruvian clade bar one (ERR7339051) which had *mmpL5* Arg202Leu. This event of reading-frame restoration is likely explained by a recent secondary duplication of a T downstream of the initial deletion (777876 GGCAT > GGAT, GGAT > GGATT). We considered the phenotype of this strain as unknown. No other *mmpL5* mutations were found in any isolate containing *mmpR5* mutations within this study though we did identify a low prevalence of variants in *mmpL5* and *mmpS5* independent of *mmpR5* mutations across both lineages (**Supplementary Figures S13 and S14**).

We also noted a tendency for *mmpR5* mutations to emerge in clades that also displayed genetic markers of rifampicin resistance. This was more common in mutations emerging after 2007 (77.2%) than before 2007 (58.3%). Most of the oldest Ile67fs Peruvian clade was rifampicin resistant (58/66 samples), with the remaining samples demonstrating only isoniazid resistance.

### Phenotypic validation of *mmpR5* variants

Given documented epistasis as a modulator of bedaquiline resistance phenotype, we performed MIC testing on a selection of available isolates and identified further MICs that have been recently published as part of the Cryptic consortium using microtitre plates (**Supplementary Table S7**)^52,53^. The epidemiological cut-off (ECOFF, defined as MIC of 95-99% of wild-type isolates) for bedaquiline has been proposed to be 0.12 or 0.25 μg/mL depending on the method used, although the final decision was to use an ECOFF of 0.25 μg/mL^53^.

We were able to identify 30 L4 isolates from Peru (including the aforementioned Peruvian clade) for MIC testing, and a further 9 MICs for L4 that had recently been published by the Cryptic consortium^52^. For the oldest dated *mmpR5* mutation emergence – the L4 Ile67fs mutation in Peruvian isolates with an associated MRCA estimated to 1701 - 10/11 (90.9%) had an MIC below the lower proposed ECOFF of 0.12 μg/mL, presumably due to the co-existing *mmpL5* loss of function mutation. Hence, we denote isolates from this clade as having a hypersusceptible phenotype. The second oldest predicted resistance mutation (Arg90Cys, dated to 1940) was however associated with MICs ≥0.12 μg/mL in 6/7 (85.7%) instances, and in 3/4 (75%) instances for the third oldest predicted resistance associated mutation for which data were available (Asn98Asp, dated to 1987). These MICs are above the wild-type range, if not formally classified as resistant. Clades with associated MIC confirmation are highlighted in Figure 3b.

## Discussion

Our work establishes that the emergence of variants in *mmpR5*, including bedaquiline RAVs, is not solely driven by the use of bedaquiline. We identified up to 11 cases where RAVs have emerged prior to the first clinical trials of bedaquiline in 2007 and a further four cases of variants emerging prior to the clinical use of bedaquiline which are expected to give rise to an intermediate phenotype. These are highlighted red and orange respectively in Supplementary Table S7, not including the oldest emergence of Ile67fs as its resistant phenotype is negated by the epistatic interaction. Phylogenetic inference estimated the oldest clade containing *mmpR5* mutations, composed mostly of samples from Peru carrying the Ile67fs RAV, to have emerged around 1702 (1657-1732). We identify two further early emergences of *mmpR5* mutations, estimated to 1871 and 1940 (Asp141fs and Arg90Cys; point estimates), with samples from the latter clade confirmed to have MICs above the wild-type range justifying classification of an intermediate phenotype. The phenotypic implications of Asp141fs remains unclear. However, together this suggests the likely circulation of variants exhibiting borderline resistance even prior to the first clinical trials for clofazimine. Our phylogenetic inference method, which points to multiple emergences of *mmpR5* nonsynonymous/frameshift variants predating the use of bedaquiline, is also confirmed by the direct observation of eight *Mtb* genomes carrying *mmpR5* RAVs sampled prior to 2007. We also identified, within the aforementioned Peruvian clade, a consistent frameshift mutation in *mmpL5*, which seemed to counteract the resistance phenotypic through an epistatic interaction (MIC <0.12 μg/mL). While Ile67fs is central for bedaquiline resistance in *Mtb*, and this mutation has clearly emerged well prior to the use of bedaquiline and clofazimine in this clade, its phenotypic impact is influenced by the strain genetic background.

We identified other localised clusters with *mmpR5* mutations, reinforcing the need for concern even in situations where such mutations are globally rare. This included 19 isolates with a Met146Thr mutation found in lineage 4 isolates from Eswatini. Met146Thr mutations have been previously associated with a clade that has a rifampicin-resistance conferring mutation located outside of the canonical rifampicin-resistance determining region, and these isolates exhibit elevated bedaquiline MICs^54^. The emergence of the Met146Thr mutation has previously been dated to have emerged in approximately 2003^23,39,54^. This is in reasonable agreement with our analysis on a much larger dataset which inferred an emergence in 2005.6 (95% confidence intervals 2004.8 – 2006.0). The long-standing presence of variants implicated in resistance and borderline resistance to bedaquiline predating the use of the drug and at high prevalence in geographically notable cases is of concern, as it suggests that non-synonymous mutations in *mmpR5* exert little fitness cost.

Together, our work suggests the existence of pre-existent reservoirs of bedaquiline resistant *Mtb*. These may have been selected for through historic clofazimine use, though we note at least one case of intermediate resistance to bedaquiline emerging as early as 1930-1947. We also note that detected variants in mmpR5 tend to exist in strains already displaying rifampicin-resistance, although also include otherwise fully susceptible strains (**Supplementary Table S7**). Together this suggests the important role of prior drug exposure in selecting for strains with pre-existing (cross-)resistance potential. This reservoir of putatively adaptive variants is expected to expand under drug pressure with the increasing use of bedaquiline and clofazimine in TB treatment. Further, these reservoirs may also pose a threat for other candidate TB agents from different drug classes that are also exported by *mmpS5* and *mmpL5*^19,22,55^.

The identification of resistance variants occurring before the clinical use of a drug is not limited to *M. tuberculosi*s and *mmpR5* alone. To illustrate, within *M. bovis*, there is evidence indicating that the *pncA* H57D mutation, which is associated with resistance to pyrazinamide (PZA), emerged approximately 900 years ago, providing inherent resistance to PZA in *M. bovis*^56^. Similarly, variations in intrinsic susceptibility to pretomanid have been observed across the MTBC, including *Mtb* lineages, even without prior exposure to nitroimidazoles^57^. It is likely that there are numerous other instances of such loss of function mutations with minimal or no impact on fitness, similar to the case of *mmpR5*. Furthermore, the existence of antimicrobial resistance (AMR) in different forms has persisted throughout the natural history of various bacteria^58^.

Nevertheless, it is crucial to determine the age and diversity of variants that have been implicated in drug resistance to gain a better understanding of the potential for widespread resistance as a contemporary challenge. We identified a large number of different *mmpR5* nonsynonymous/frameshift variants across both of our *Mtb* lineage cohorts; 46 in L2 and 67 in L4. This suggests the mutational target leading to bedaquiline resistance is wider than for most other current TB drugs and raises concerns about the ease with which bedaquiline resistance can emerge during treatment. It is further concerning that resistance to the new class of nitroimidazole drugs, such as pretomanid and delamanid, is also conferred by loss of function mutations in any of at least six genes, suggesting that they may also have a low barrier to resistance^59^.

While we identified many non-synonymous variants in *mmpR5*, only one (Ile67fs) has been previously definitively linked to resistance. We acknowledge that several of our detected variants have no associated MIC values available in the literature and are thus currently not fully phenotypically validated. We hope by presenting these as ‘unknown’ our work, estimating the age of emergence of non-synonymous mutations, can be of value as further variants are phenotyped in the future. It is however true that determining the phenotypic consequences of *mmpR5* variants that have previously been described is challenging as there are often only limited reports correlating MICs to genotypes. Moreover, at least four different methods are used to determine MICs, some of which do not have associated critical concentrations. Even where critical concentrations have been set, there is an overlap in MICs of isolates that are genetically wild type and those that have mutations likely to cause resistance^35^. We also note that as we purposefully enriched our dataset for *mmpR5* mutations, the sampling precludes estimation of the overall prevalence of these mutations in genome sequencing databases.

Prediction of phenotypic bedaquiline resistance from genomic data is further complicated by the existence of hypersusceptibility variants. For example, the c-11a variant located in the promoter of mmpR5, which appears to increase susceptibly to bedaquiline^38^, was observed to be fixed throughout a large clade within L2. The early emergence of this variant and its geographical concentration in South Africa and Eswatini may suggest the role of non-pharmacological influences on mmpR5 which regulates multiple MmpL efflux systems^20^. Further, analysis of hypersusceptibility is limited by the truncated lower MIC range of the UKMYC microtitre plates, with many isolates giving MICs below the lower end of the measured range. While large-scale genotype/phenotype analyses will likely support the development of rapid molecular diagnostics, targeted or whole genome sequencing, at reasonable depths, may provide the only opportunity to detect all possible *mmpR5* RAVs, and possible co-occurring mutations, in clinical settings.

Bedaquiline resistance can also be conferred by other RAVs including in *pepQ* (bedaquiline and clofazimine), *atpE* (bedaquiline only)^51^ and *Rv1979c* (clofazimine only). We only found *atpE* RAVs at low allele frequency in one patient who also had *mmpR5* variants (sample accession ERR2677436), which is in line with other evidence suggesting they rarely occur in clinical isolates, likely due to a high fitness cost. Likewise, we only identified *Rv1979c* RAVs in three patients in China, although there were other variants in *Rv1979c* for which ability to cause phenotypic resistance has not been previously assessed. Frameshift *pepQ* mutations that are potentially causative of resistance were identified in 11 cases, in keeping with its possible role as an additional rare resistance mechanism.

Our findings, of reservoirs of *mmpR5* RAVs predating the therapeutic use of bedaquiline, are of high clinical relevance as the presence of *mmpR5* variants during therapy in clinical strains has been associated with substantially worse outcomes in patients treated with drug regimens including bedaquiline^36^. Although it is uncertain what the impact of *mmpR5* RAVs are on outcomes when present prior to treatment^60,61^, it is imperative to monitor and prevent the wider transmission of bedaquiline resistant clones, particularly in high MDR/XDR-TB settings. Early evaluation of new TB drug candidates entering clinical trials will also be vital given early data suggesting possible cross-resistance for DprE1 inhibitors such as macozinone^22^. The large and disparate set of mutations in *mmpR5* we identified, with differing phenotypes and some having been in circulation historically, adds further urgency to the development of rapid drug susceptibility testing for bedaquiline to inform effective treatment choices and mitigate the further spread of DR-TB.

## Materials and methods

### Sample collection

In this study we curated large representative datasets of *Mtb* whole genome sequences encompassing the global genetic and geographic distribution of lineages 2 (L2) and L4 (**Figure 1, Supplementary Tables S1-S2**). The dataset was enriched to include all available sequenced isolates with *mmpR5* variants, which in some cases included isolates with no, or limited, published metadata. In all other cases samples for which metadata on the geographic location and date of collection was available were retained. To ensure high quality consensus alignments we required that all samples mapped with a minimum percentage cover of 96% and a mean coverage of 30x to the H37Rv reference genome (NC_000962.3). We excluded any samples with evidence of mixed strain infection as identified by the presence of lineage-specific SNPs to more than one sublineage^62^ or the presence of a high proportion of heterozygous alleles^63^. The total number of samples included in these datasets, and their source is shown in **Supplementary Table S2.** An index of all samples is available in **Supplementary Table S1**.

A large global dataset of 1,669 L4 *Mtb* sequences has been constructed, which we used as the basis for curating our L4 dataset^44^. We refer to this as the ‘base dataset’ for L4. For L2, we constructed a ‘base dataset’ by screening the Sequence Read Archive (SRA) and European Nucleotide Archive (ENA) using BIGSI^64^ for the *rpsA* gene sequence containing the L2 defining variant *rpsA* a636c^62^ with a 100% match. This search returned 6,307 *Mtb* genomes, of which 1,272 represented unique samples that had the minimum required metadata. Metadata from three studies were also added manually as they were not included in their respective SRA submissions but were available within published studies^65–67^.

For isolates with only information on the year of sample collection, we set the date to be equal to the middle of the year. For those with information on the month but not the date of collection we set the date of collection to the first of the month. For sequenced samples which were missing associated metadata (32 L2 genomes and 19 L4 genomes) we attempted to estimate an average time of sample collection in order to impute a sampling date. To do so we computed the average time between date of collection and sequence upload date for all samples with associated dates separately in each of the L2 and L4 datasets (**Supplementary Figure S1**). For L2 we estimated a mean lag time of 4.7 years (0.5– 12.6 years 95% CI). For L4, having excluded three sequences obtained from 18^th^ Century mummies from Hungary^40^, we estimated a mean lag time of 6.9 years (0.6-19.1 years 95% CI). The estimated dates, where required, are provided in Supplementary Table S1.

To enrich the datasets for isolates with *mmpR5* variants, we included further sequences from our own studies in KwaZulu-Natal, South Africa^41,42^, other studies of drug-resistant TB in southern Africa^16,44,68–71^, and Peru^72,73^. We additionally supplement the Peruvian data with 163 previously unpublished isolates. In these cases, and to facilitate the most accurate possible estimation of the date of resistance emergence, we included samples with *mmpR5* variants as well as genetically related sequences without *mmpR5* variants.

To identify further published raw sequencing data with *mmpR5* variants from studies where bedaquiline/clofazimine resistance may have been previously unidentified, we screened the NCBI Sequencing Read Archive (SRA) for sequence data containing 85 previously published *mmpR5* variants^28–30,41,42,74,75^ with BIGSI^64^. BIGSI was employed against a publicly available indexed database of complete SRA/ENA bacterial and viral whole genome sequences current to December 2016 (available here: http://ftp.ebi.ac.uk/pub/software/bigsi/nat_biotech_2018/all-microbial-index-v03/), and also employed locally against an updated in-house database which additionally indexed SRA samples from January 2017 until January 2019. Samples added using this approach are flagged ‘BIGSI’ in **Supplementary Table S1**. We also used the PYGSI tool^76^ to interrogate BIGSI with the *mmpR5* sequence adjusted to include every possible single nucleotide substitution. In each instance we included 30 bases upstream and downstream of the gene as annotated on the H37Rv *Mtb* reference genome. For the purpose of this study we only considered coding region, non-synonymous substitutions and insertions and deletions. Samples added following the PYGSI screen are flagged ‘PYGSI’ in **Supplementary Table S1**. A breakdown of the different datasets used is provided in **Supplementary Table S2**.

### Reference mapping and variant calling

Original fastq files for all included sequences were downloaded and paired reads mapped to the H37Rv reference genome with bwa mem v0.7.17^77^. Mapped reads were sorted and de-duplicated using Picard Tools v2.20 followed by indel realignment with GATK v3.8^78^. Alignment quality and coverage was recorded with Qualimap v2.21^79^. Variant calling was performed using bcftools v1.9, based on reads mapping with a minimum mapping quality of 20, base quality of 20, no evidence of strand or position bias, a minimum coverage depth of 10 reads, and a minimum of four reads supporting the alternate allele, with at least two of them on each strand. Moreover, SNPs that were less than 2bp away from an indel were excluded from the analysis. Similarly, only indels 3bp apart of other indels were kept.

All sites with insufficient coverage to identify a site as variant or reference were excluded (marked as ‘N’), as were those in or within 100 bases of PE/PPE genes, or in insertion sequences or phages. SNPs present in the alignment with at least 90% frequency were used to generate a pseudoalignment of equal length to the H37Rv. Samples with more than 10% of the alignment represented by ambiguous bases were excluded. Those positions with more than 10% of ambiguous bases across all the samples were also removed. In order to avoid bias on the tree structure, positions known to be associated with drug resistance were not included.

A more permissive variant calling pipeline was used to identify *mmpR5* variants, as they are often present at <100% frequency with a high incidence of frameshift mutations. Here we instead employed FreeBayes v1.2^85^ to call all variants present in the *mmpR5* gene (or up to 100 bases upstream) that were present at ≥5% frequency (alternate allele fraction *-F* 0.05) and supported by at least four reads including one on each strand. Using this more permissive variant calling strategy we also systematically screened for all mutations in the efflux pump proteins mmpS5-mmpL5 operon (**Supplementary Figures S13 and S14**).

### Classification of resistance variants

All raw fastq files were screened using the rapid resistance profiling tool TBProfiler^47,80^ against a curated whole genome drug resistance mutations library. This allowed rapid assignment of polymorphisms associated with resistance to different antimycobacterial drugs and categorisation of MDR and XDR *Mtb* status (**Supplementary Figure S2, Supplementary Figures S5-S9**). Resistance profiles of sequences containing *mmpR5* variants are listed in Supplementary Table S7 as either “S” for susceptible, “RR” for rifampicin-resistant and “preXDR” for fluoroquinolone-resistant.

### Classification of *mmpR5* variants

The diverse range of *mmpR5* variants and paucity of widespread MIC testing means that there are limited data from which to infer the phenotypic consequences of identified *mmpR5* variants. This was true aside from a subset of data sampled in Peru for which 30 L4 isolates from Peru were subjected to MIC testing using the UKMYC6 plate and a further nine were evaluated for MICs reported by the Cryptic consortium^52^. The approach we used was to assign whether nonsynonymous variants confer a normal or raised MIC based on published phenotypic tests for strains carrying that variant. A full list of the literature reports used for each mutation is provided in **Supplementary Table S4**. We also introduced an intermediate category to describe isolates with MICs at the critical concentration (e.g., 0.25µg/mL on Middlebrook 7H11 agar), where there is an overlap of the MIC distributions of *mmpR58* mutated and wild type isolates with uncertain clinical implications^35^. We assumed that all other disruptive frameshift and stop mutations would confer resistance in light of the role of *mmpR5* as a negative repressor, where loss of function should lead to efflux pump overexpression, unless evidence exists in the literature to suggest otherwise. This allowed us to identify two frameshifts of currently unclear effect (**Supplementary Table S4**). All other promoter and previously unreported missense mutations were categorised as unknown (**Supplementary Table S4**). Where *mmpR5* mutations were accompanied by an *mmpS5* or *mmpL5* loss of function mutation, we assumed that would confer susceptibility (or hypersusceptibility) to bedaquiline^23^.

### Global phylogenetic inference

The alignments for phylogenetic inference were masked for the *mmpR5* region using bedtools v2.25.0. All variant positions were extracted from the resulting global phylogenetic alignments using snp-sites v2.4.1^81^, including a L4 outgroup for the L2 alignment (NC_000962.3) and a lineage 3 (L3) outgroup for the L4 alignment (SRR1188186). This resulted in a 67,585 SNP alignment for the L4 dataset and 29,205 SNP alignment for the L2 dataset. A maximum likelihood phylogenetic tree was constructed for both SNP alignments using RAxML-NG v0.9.0^82^ specifying a GTR+G substitution model, correcting for the number of invariant sites using the ascertainment flag (ASC_STAM) and specifying a minimum branch length of 1×10^−9^ reporting 12 decimal places (--precision 12).

### Estimating the age of emergence of *mmpR5* variants

To test whether the resulting phylogenies can be time-calibrated we first dropped the outgroups from the phylogeny and rescaled the trees so that branches were measured in unit of substitutions per genome. We then computed a linear regression between root-to-tip distance and the time of sample collection using BactDating^83^, which additionally assesses the significance of the regression based on 10,000 date randomisations. We obtained a significant temporal correlation for both the L2 and L4 phylogenies, both with and without imputation of dates for samples with missing metadata (**Supplementary Figure 3**).

We employed the Bayesian method BactDating v1.01^83^, run without updating the root (updateRoot=F), a mixed relaxed gamma clock model and otherwise default parameters to both global datasets. The MCMC chain was run for 1×10^7^ iterations and 3×10^7^ iterations. BactDating results were considered only when MCMC chains converged with an Effective Sample Space (ESS) of at least 100. The analysis was applied to the datasets both with and without considering imputed and non-imputed collection dates (**Supplementary Table 3**).

To independently infer the evolutionary rates associated with each of our datasets, we sub-sampled both the L4 and L2 datasets to 200 isolates, selected so as to retain the maximal diversity of the tree using Treemmer v0.3^84^. As before, we excluded all variants currently implicated in drug resistance from the alignments. This resulted in a dataset for L4 comprising 25,104 SNPs and spanning 232 years of sampling and for L2 comprising 8,221 SNPs and spanning 24 years of sampling. In both cases the L3 sample SRR1188186 was used as an out-group given this has an associated collection date. Maximum likelihood trees were constructed using RaXML-NG v0.9.0^82^, as previously described, and a significant temporal regression was obtained for both sub-sampled datasets (**Supplementary Figure S4**).

BEAST2 v2.6.0^48^ was run on both subsampled SNP alignments allowing for model averaging over possible choices of substitution models^85^. All models were run with either a relaxed or a strict prior on the evolutionary clock rate for three possible coalescent demographic models: exponential, constant and skyline. To speed up the convergence, the prior on the evolutionary clock rate was given as a uniform distribution (limits 0 to 10) with a starting value set to 10^−7^. In each case, the MCMC chain was run for 500,000,000 iterations, with the first 10% discarded as burn-in and sampling trees every 10,000 chains. The convergence of the chain was inspected in Tracer 1.7 and through consideration of the ESS for all parameters (ESS>200). The best-fit model to the data for these runs was assessed through a path sampling analysis^86^ specifying 100 steps, 4 million generations per step, alpha = 0.3, pre-burn-in = 1 million generations, burn-in for each step = 40%. For both datasets, the best supported strict clock model was a coalescent Bayesian skyline analysis. The rates (mean and 95% HPD) estimated under these subsampled analyses (L2 7.7×10^−8^ [4.9×10^−8^ - 1.03×10^−7^] substitutions per site per year; L4 7.1×10^−8^ [6.2×10^−8^ - 7.9×10^−8^] substitutions per site per year) were used to rescale the maximum likelihood phylogenetic trees generated across the entire L2 and L4 datasets, by transforming all branch lengths of the tree from per unit substitution to per unit substitutions per site per year using the R package Ape v5.3^87^. This resulted in an estimated tMRCA of 1332CE (945CE-1503CE) for L2 and 853CE (685CE – 967CE) for L4 (**Figure 2**).

The resulting phylogenetic trees were visualised and annotated for place of geographic sampling and *mmpR5* variant status using ggtree v1.14.6^88^. All nonsynonymous/frameshift mutations in *mmpR5* were considered, with the phenotypic status assigned in **Supplementary Table S4**. For the purpose of this analysis, and to be conservative, ‘unknown’ variants classified using XGBoost were still considered ‘unknown’ (**Supplementary Note 1**). Clades carrying shared variants in *mmpR5* were identified and the distributions around the age of the node (point estimates – mean - and 95% HPDs) were extracted from the time-stamped phylogeny. For isolated samples (single emergences) exhibiting variants in *mmpR5*, the time of sample collection was extracted together with the date associated with the upper bound on the age of the next closest node of the tree, allowing for the mutation to have occurred anywhere over the length of the terminal branch (**Figure 3, Supplementary Figures S11-S12**). For the Peruvian clade Bayesian skyline analysis was implemented through the skylineplot analysis functionality available in Ape v5.3^87^.

### Data availability

Raw sequence data and full metadata for all newly generated isolates are available on NCBI Sequencing Read Archive under BioProject ID: PRJEB39837.

## Author Contributions

LvD, CN and FB conceived and designed the study. JM, NP, AG, MO, AP, OBB, VE and LG provided sequence data. ATO, JP, MA, CCST and XD performed and advised on computational analyses. LvD, CN and FB wrote the manuscript with input from all co-authors. All authors read and approved the final manuscript.

## Acknowledgments

CN and JM are supported by the Wellcome Trust (203583/Z/16/Z and 203919/Z/16/Z, respectively). LvD is supported by a UCL Excellence Fellowship. F.B. acknowledges support from the BBSRC (equipment grant BB/R01356X/1). FB additionally acknowledges the National Institute for Health Research University College London Hospitals Biomedical Research Centre. M.A. was supported by a Ph.D. scholarship from University College London. All authors acknowledge UCL Biosciences Big Data equipment grant from BBSRC (BB/R01356X/1).

