## Supplement for "Detection of a historic reservoir of bedaquiline / clofazimine resistance associated variants in *Mycobacterium tuberculosis*"

**Supplementary Figures**
Pages 2-14

**Supplementary Tables**
Pages 15-21
Supplementary Table S1, S7 and S8 are available as external excel spreadsheets.

**Supplementary Note**
Pages 22-26

**References**
Pages 27-28

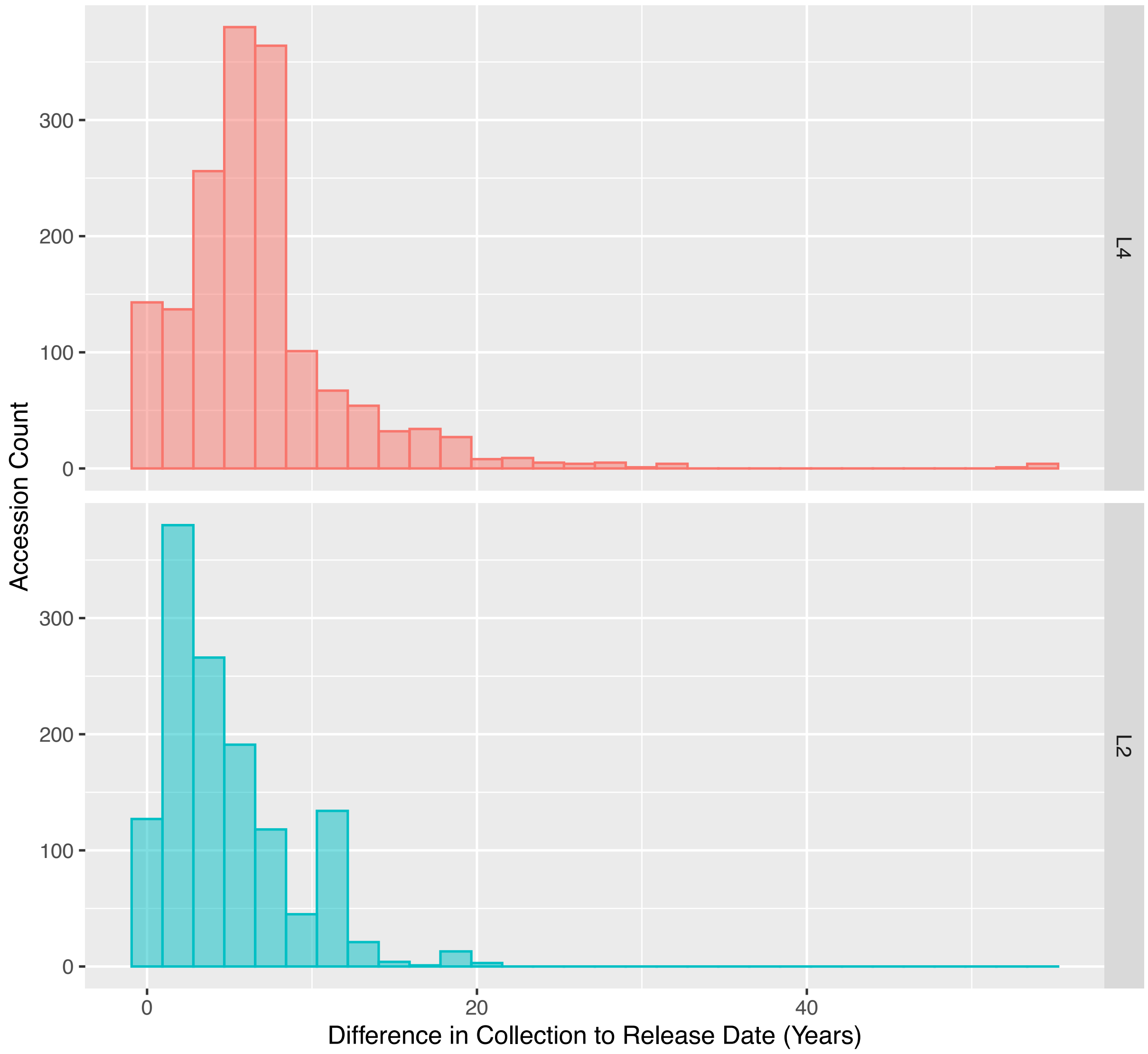


**Figure S1:** Distribution of difference (in decimal years) of collection date to release date for sequences in the lineage 2 (L2) and lineage 4 (L4) dataset. For lineage 4, having excluded samples from three $18^{th}$ century mummies, an average of 6.9 years (0.6-19.1 years 95% CI) was obtained. For lineage 2, an average lag-time of 4.7 years (0.5–12.6 years 95% CI) was obtained.

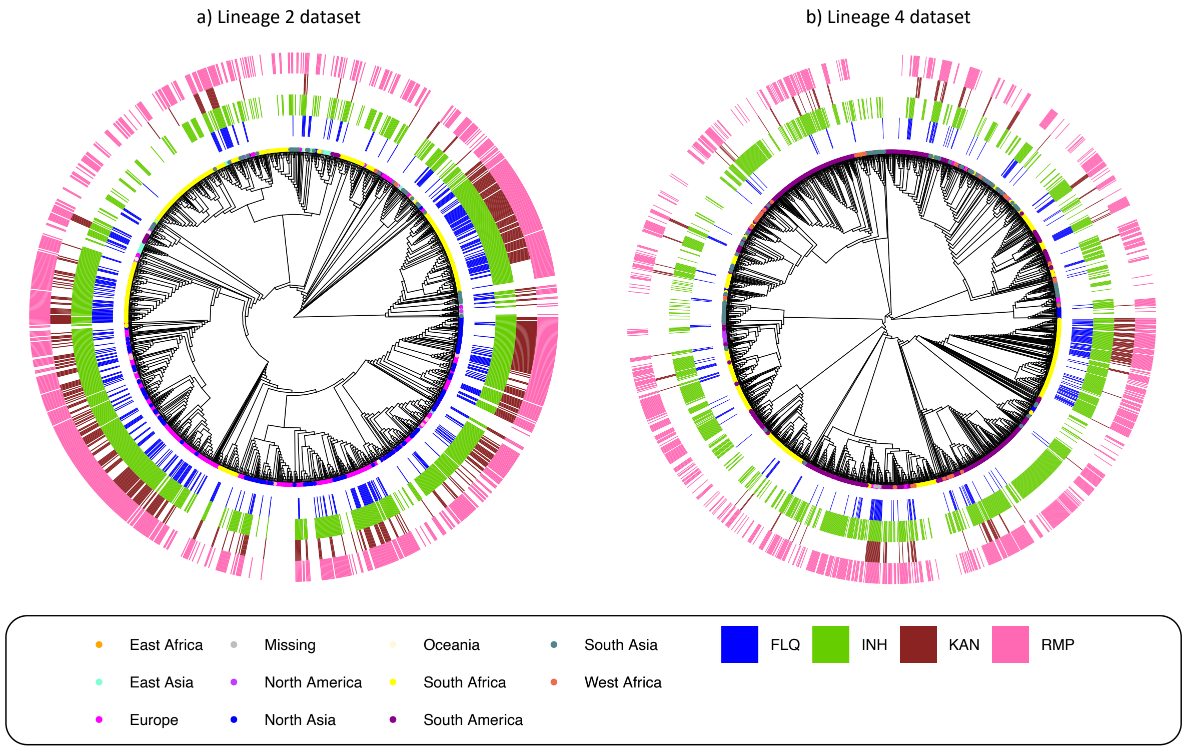


**Supplementary Figure S2:** Maximum likelihood phylogenetic tree of the lineage 2 (a) and lineage 4 (b) datasets. Tip colours provide the country of sample collection and outer bars give the resistance to four antimycobacterial drugs: fluroquinolones (FLQ), isoniazid (INH), kanamycin (KAN) and rifampicin (RMP).

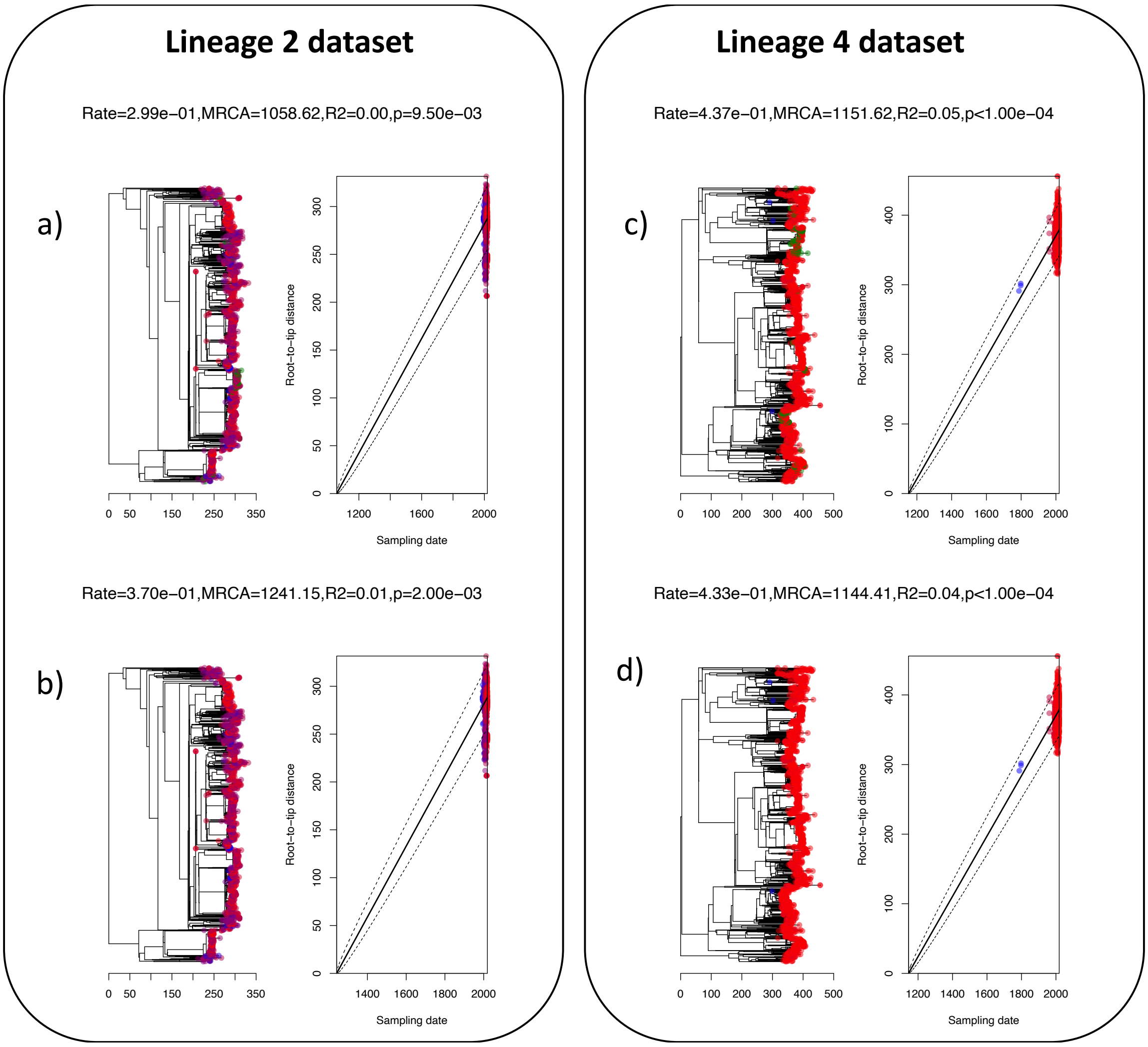


**Supplementary Figure S3:** Linear regressions of root-to-tip distance (y-axis) versus sampling dates (x-axis) for global *Mtb* datasets; lineage 2 (a-b) and lineage 4 (c-d). Regressions are performed both without (a-c) and with (b-d) imputation of missing dates (see Methods). Here, the *p*-value (tip-randomisation test) is calculated by fitting a linear regression to the root to tip distance vs sampling date for 10,000 randomisations and adding the number of randomised fits that present a better regression coefficient than the real data (divided by 10,000).

#### a) Lineage 2

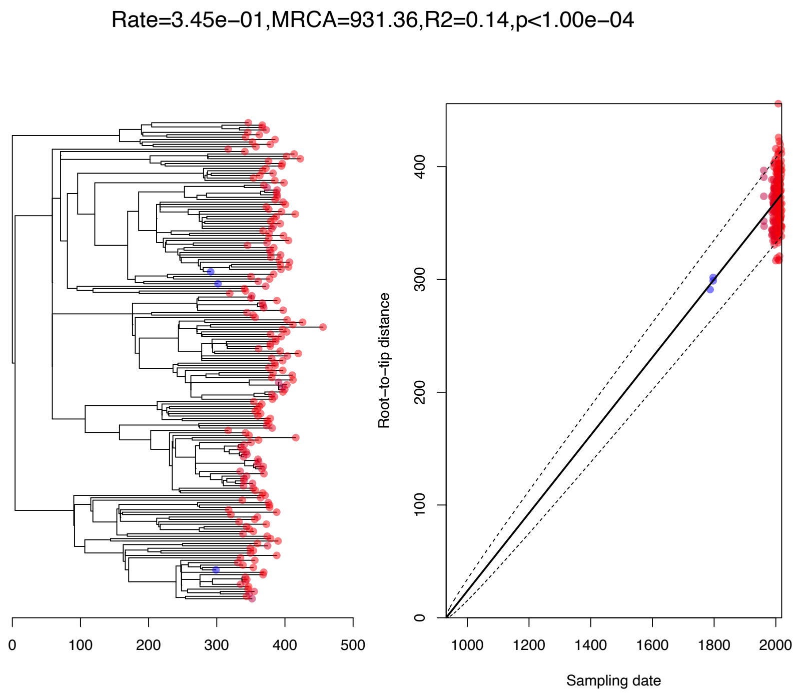


#### b) Lineage 4

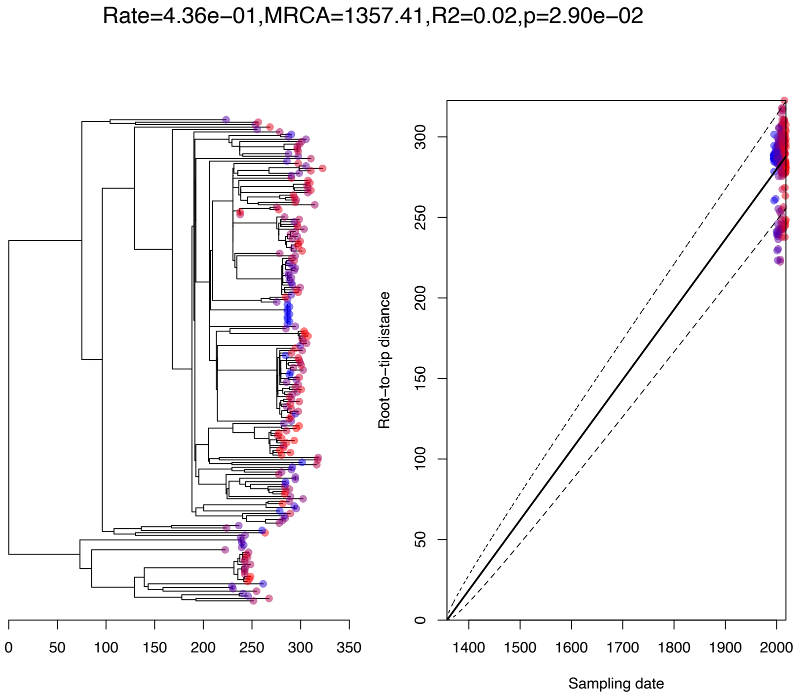


**Supplementary Figure S4:** Sub-sampled datasets temporal regression. The lineage 2 data covered 24 years of molecular evolution. The lineage 4 dataset comprised 232 years of evolution.

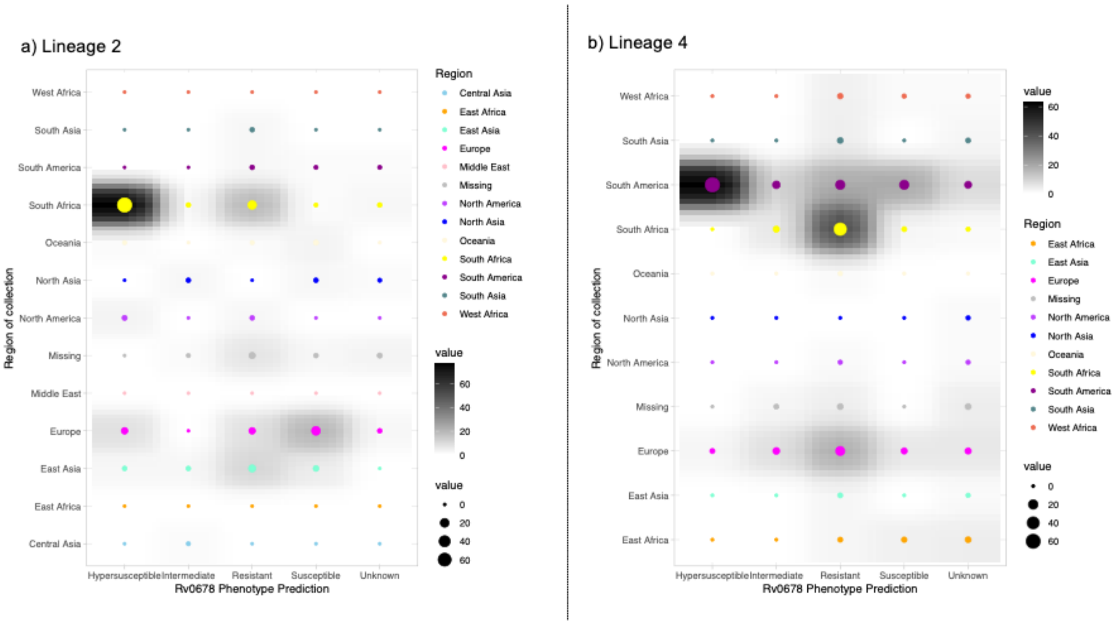


**Supplementary Figure S5:** Number (count) of *mmpR5* variants split by predicted phenotype (x-axis) for each geographic region included in each of the lineage 2 (a) and lineage 4 b) datasets.

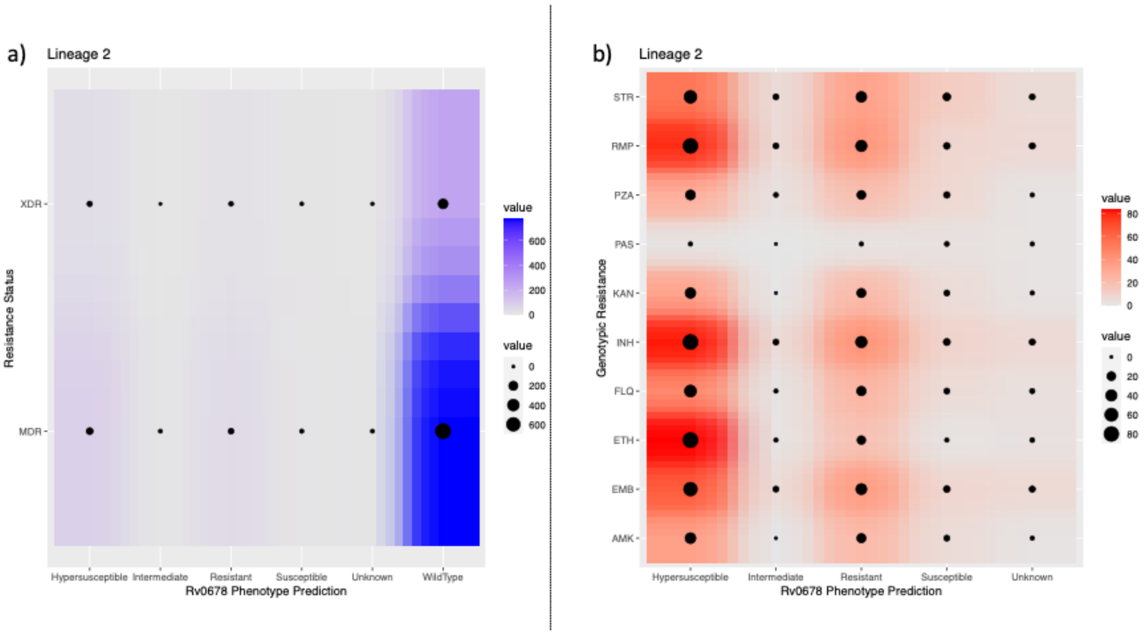


**Supplementary Figure S6:** Co-occurrence count of *mmpR5* variant phenotype predictions (x-axis) and resistance status a) and genotypic resistances b) for the lineage 2 dataset.

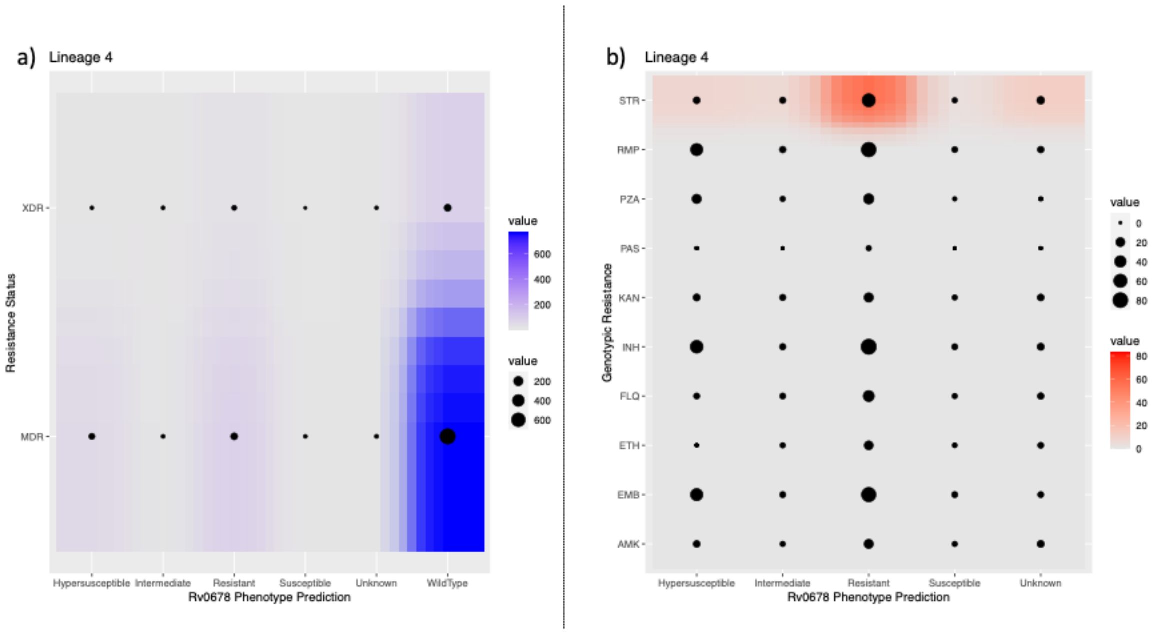


**Supplementary Figure S7:** Co-occurrence count of *mmpR5* variant phenotype predictions (x-axis) and resistance status a) and genotypic resistances b) for the lineage 4 dataset.

| Classification | MDR | S | XDR |
|---|---|---|---|
| Hypersusceptible | 48 | 7 | 30 |
| Intermediate | 5 | 3 | 0 |
| Resistant | 24 | 11 | 16 |
| Susceptible | 5 | 21 | 3 |
| Unknown | 4 | 2 | 1 |
| Wildtype | 530 | 330 | 245 |

*Chi=51.875, df=10, p=1.2e-07*

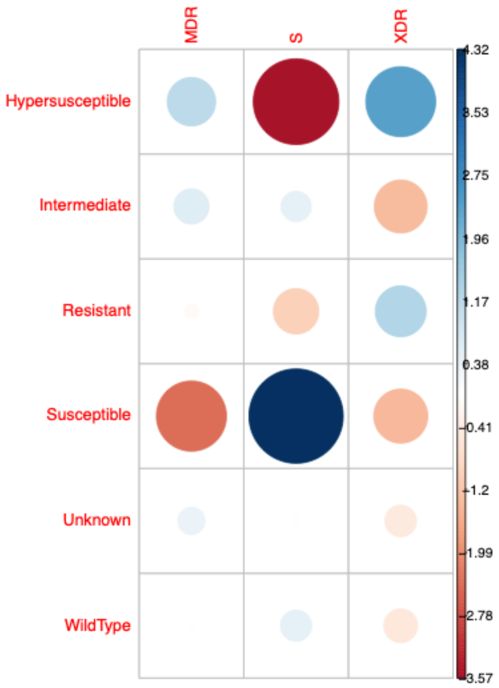


**Supplementary Figure S8:** Contingency tables and $\chi^2$ results for the distribution of susceptible, MDR and XDR *Mtb* amongst strains carrying variants in *mmpR5* in the lineage 2 dataset. Plots provide the squared standardised residuals contributing to the rejection of the null hypothesis. The colour intensity and size of the circle is proportion to the contribution with positive displayed in blue and negative in red.

| Classification | MDR | S | XDR |
|---|---|---|---|
| Hypersusceptible | 43 | 11 | 4 |
| Intermediate | 1 | 17 | 5 |
| Resistant | 57 | 11 | 19 |
| Susceptible | 3 | 25 | 2 |
| Unknown | 3 | 10 | 4 |
| Wildtype | 689 | 702 | 81 |

*Chi=127.83, df=10, p<2.2e-16*

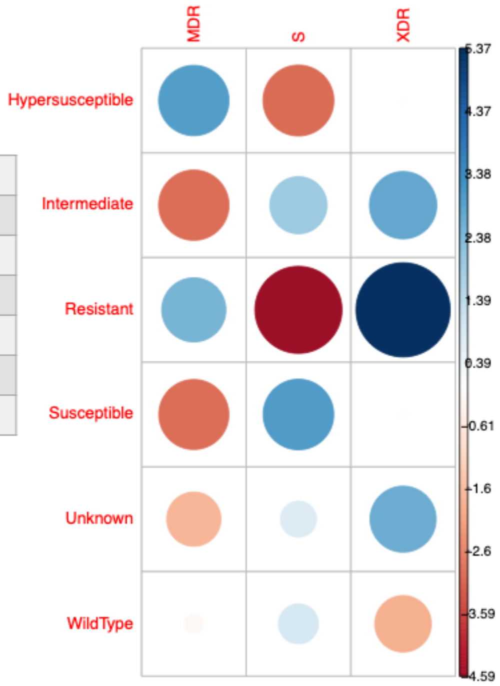


**Supplementary Figure S9:** Contingency tables and $\chi^2$ results for the distribution of susceptible, MDR and XDR *Mtb* amongst strains carrying variants in *mmpR5* in the lineage 4 dataset. Plots provide the squared standardised residuals contributing to the rejection of the null hypothesis. The colour intensity and size of the circle is proportion to the contribution with positive displayed in blue and negative in red.

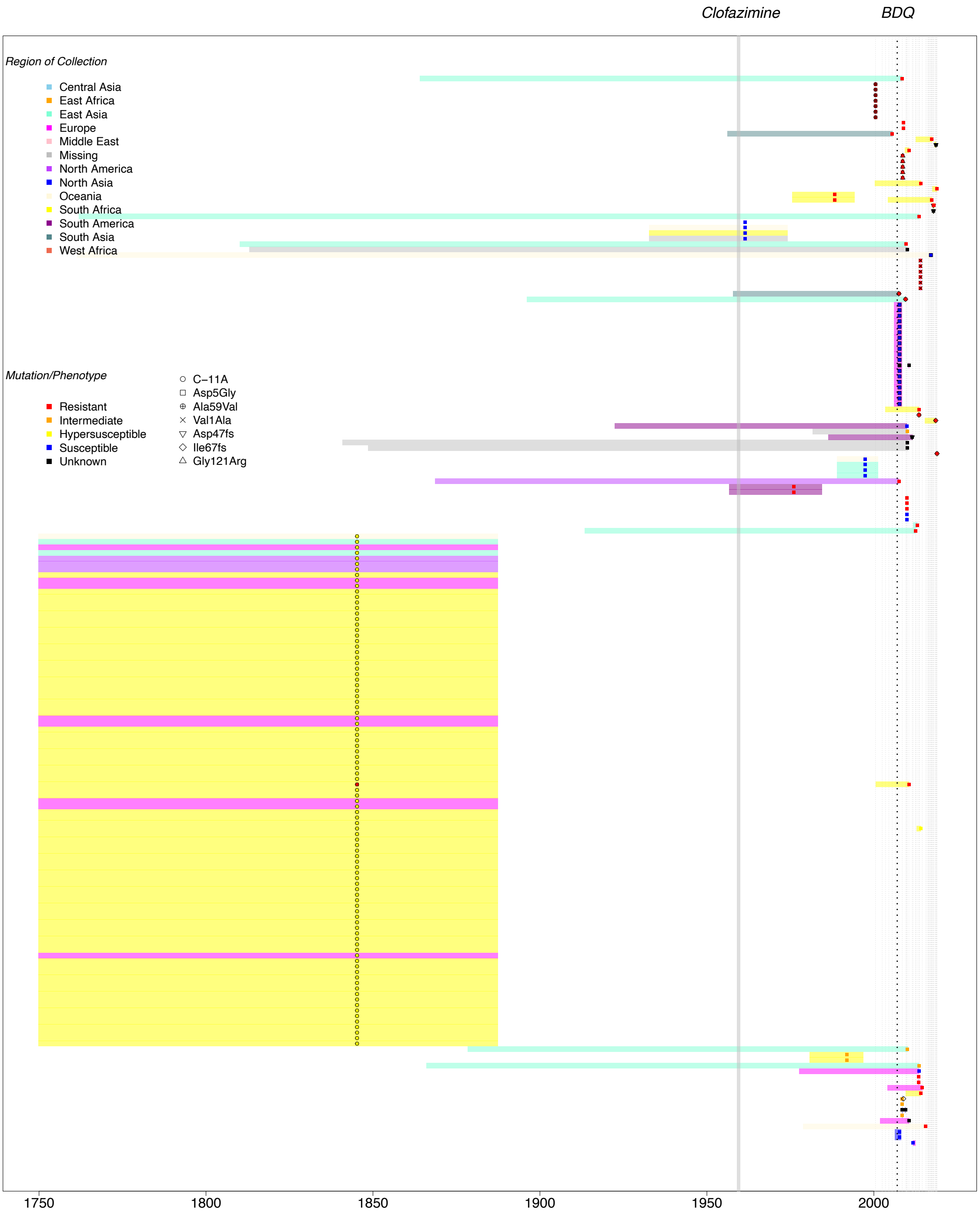


**Supplementary Figure S10:** Full mutational timeline for the estimated date of emergence (x-axis) and confidence intervals of nodes with descendent tips carrying nonsynonymous variants in *mmpR5* in the lineage 2 dataset. All nonsynonymous variants are depicted. Confidence bars are coloured according to the region where the isolate was collected. Symbols provide the point estimates of the age of the node coloured by *mmpR5* predicted phenotype. Symbols are used for all mutations occurring in ≥5 isolates, in this case. Grey dashed lines provide the collection date of all sequenced isolates included in the analysis with *mmpR5* variants. Data available in Supplementary Table S7.

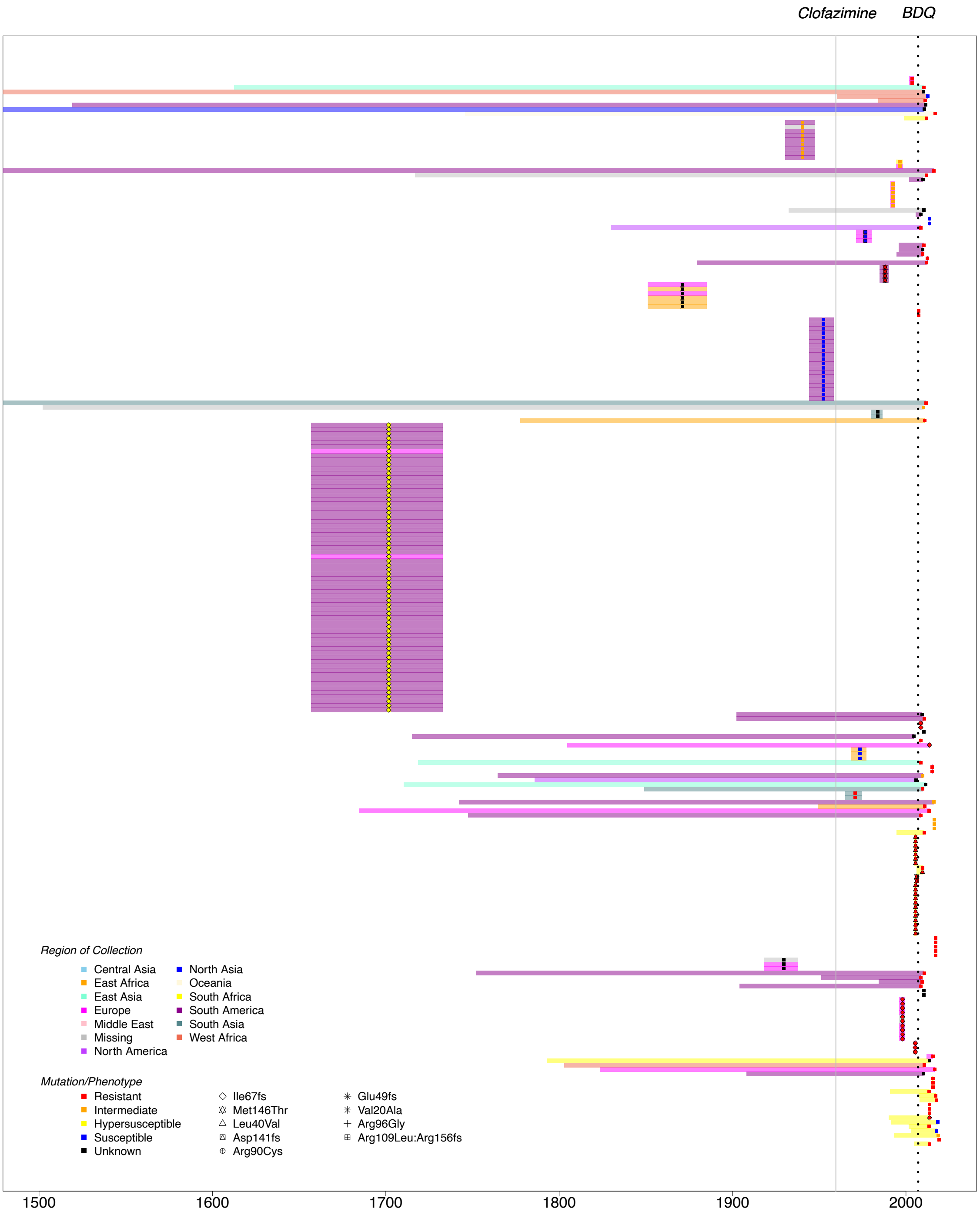


**Supplementary Figure S11:** Full mutational timeline for the estimated date of emergence (x-axis) and confidence intervals of nodes with descendent tips carrying nonsynonymous variants in *mmpR5* in the lineage 4 dataset. All nonsynonymous variants are depicted. Confidence bars are coloured according to the region where the isolate was collected. Symbols provide the point estimates of the age of the node coloured by *mmpR5* predicted phenotype. Symbols are used for all mutations occurring in ≥5 isolates. Grey dashed lines provide the collection date of all sequenced isolates included in the analysis with *mmpR5* variants. Data available in Supplementary Table S7.

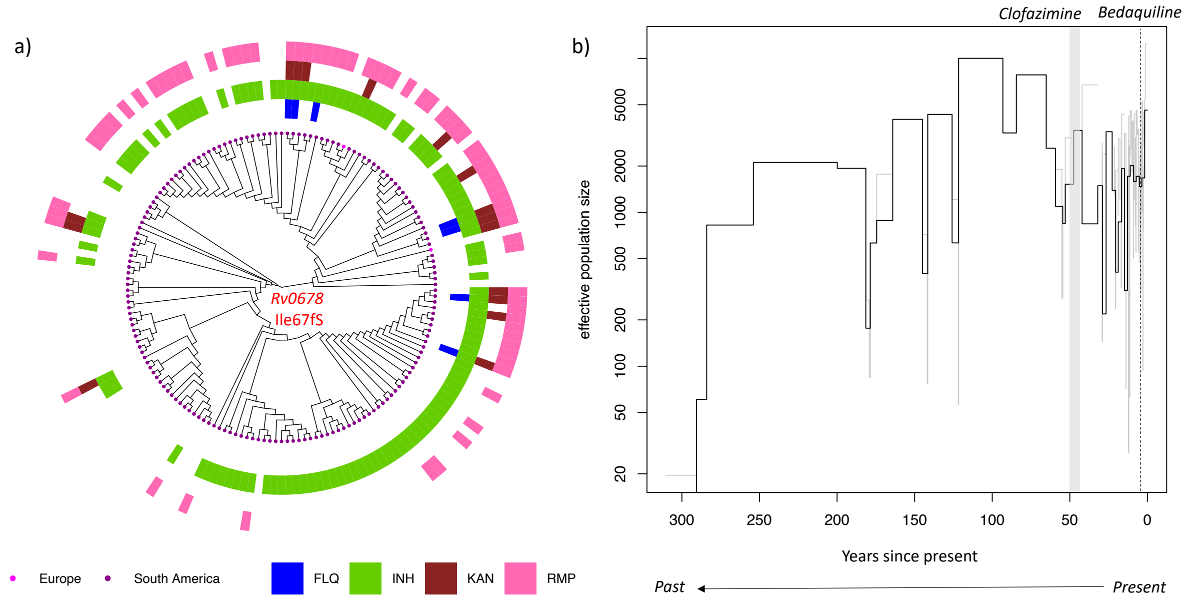


**Supplementary Figure S12**: a) Lineage 4 Peruvian *mmpR5* Ile67fs + mmpL5 Arg202fs carrying clade phylogeny, which has a tMRCA dating to 1702 (1657-1732). Phenotypic resistances for fluoroquinolones (FLQ), isoniazid (INH), kanamycin (KAN) and rifampicin (RMP) are provided as outer coloured rings. Most samples are from Peru (purple), though two samples are from Europe (Sweden and the Netherlands). b) Provides the generalized skyline plot estimate of effective population size through time based on the timed phylogeny of this clade. Grey lines provide the full skyline plot, black lines provide the coalescent intervals. The first clinical use of clofazimine and bedaquiline are provided by the axis at top.

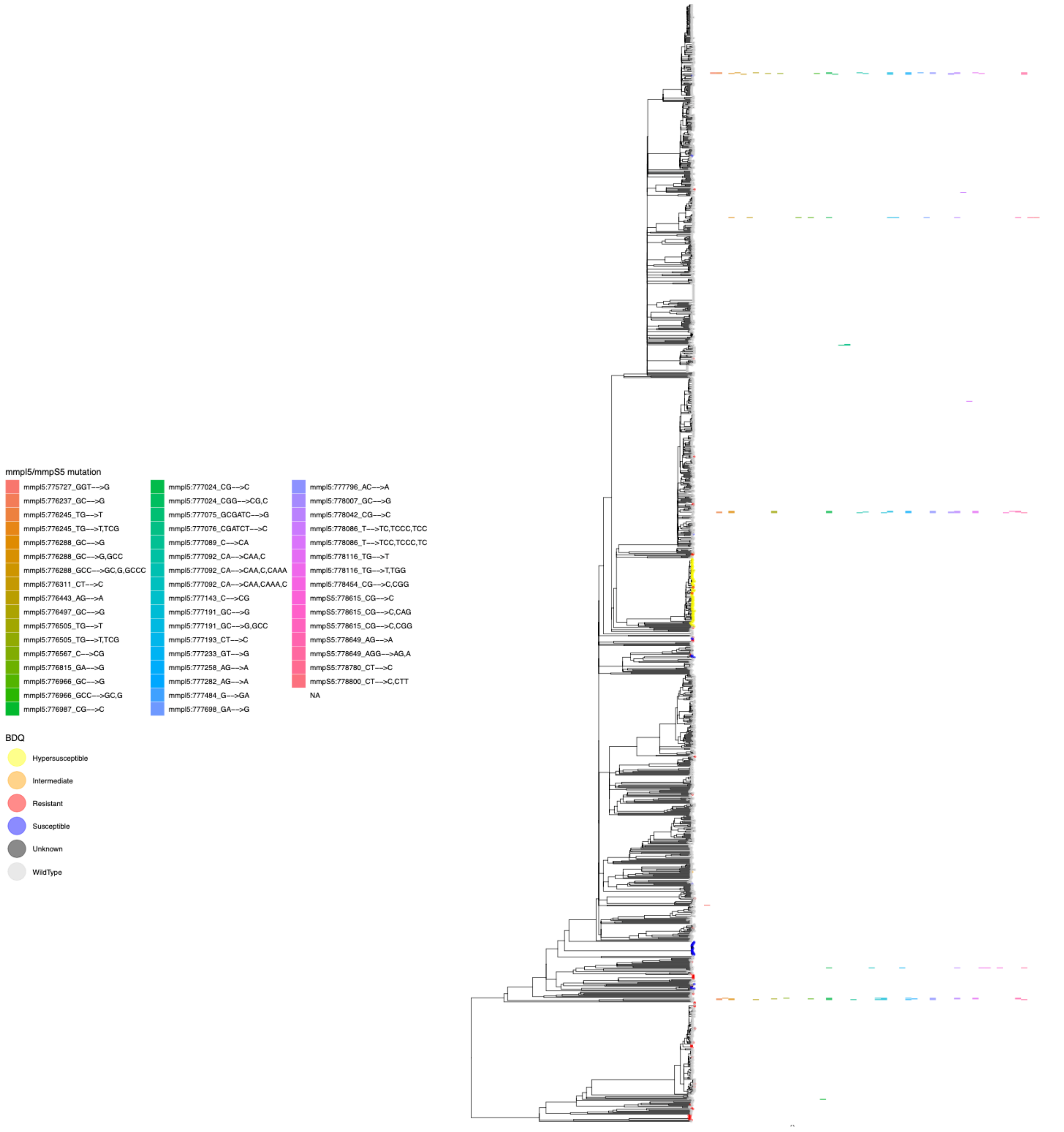


**Supplementary Figure S13**: Phylogenetic distribution of LOF mutations identified in *mmpL5* and *mmpS5* identified in L2 isolates. Phylogeny is provided with tip colours according to inferred bedaquiline resistance status. Heatmap provides colour for presence of a mutation as ordered by the vertical columns.

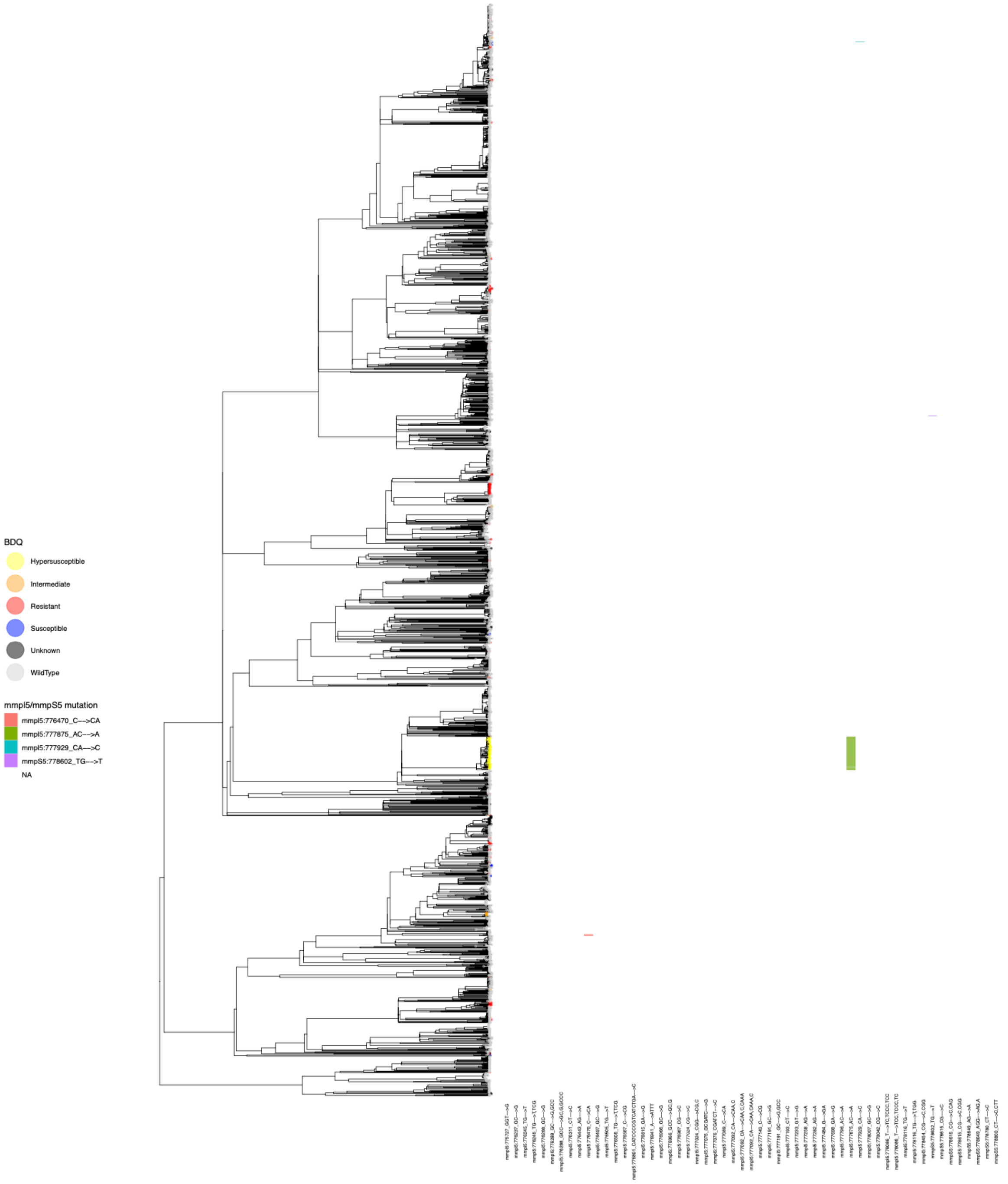


**Supplementary Figure S14**: Phylogenetic distribution of LOF mutations identified in *mmpL5* and *mmpS5* identified in L4 isolates. Phylogeny is provided with tip colours according to inferred bedaquiline resistance status. Heatmap provides colour for presence of a mutation as ordered by the vertical columns.

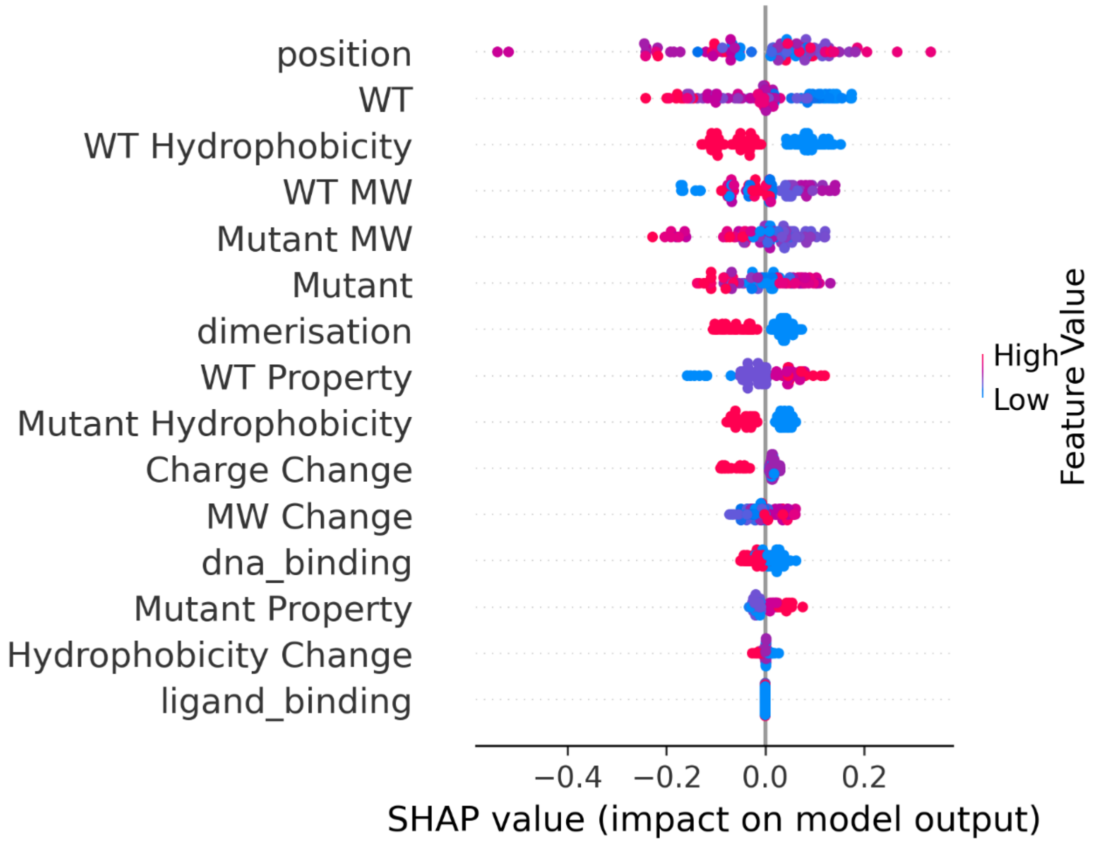


**Supplementary Figure S15:** Summary plot of SHAP values. Each point represents the SHAP value of a single prediction for a particular feature. Points are stacked vertically using density estimation. ‘WT’, ‘mutant’ and ‘MW’ denotes the wild type amino acid, amino acid variant and molecular weight (Da) respectively. ‘Property’ refers to whether an amino acid was non-polar, polar, positively charged or negatively charged. ‘Ligand_binding’, ‘dna_binding’ and ‘dimerisation’ refer to whether the amino acid residue is involved in ligand binding, DNA binding or dimerisation. ‘Position’ refers to the integer 5’-3’ position of the variant. Positive SHAP values imply an increase in the predicted probability of resistance due to the presence of the feature.

**Table S1 [external excel file]:** Spreadsheet providing the whole genomes sequences included in both alignments with accession numbers and *mmpR5* variants annotated.

**Table S2:** Source of whole genome sequences included in the global lineage 2 and lineage 4 alignments following quality checks and as given in Table S1. The number in brackets designate those with *mmpR5* differing from wild type.

| **Data source** | **Sequence count (with *mmpR5* variants)** | |
|---|---|---|
| | **Lineage 2** | **Lineage 4** |
| Base dataset | 1016 (12) | 1166 (20) |
| South Africa – our centre | 155 (11) | 243 (17) |
| Southern Africa – other studies (marked 'Additional' in Table S1) | 238 (77) | 426 (73) |
| Peru | 11 (3) | 266 (70) |
| BIGSI | 73 (173) | 40 (40) |
| PYSGI | 18 (18) | 24 (24) |
| Strain Bank sequences [1] | 3 (0) | 3 (0) |
| Total | 1514 (294) | 2168 (244) |

**Table S3:** Summary of phylogenetic dating approaches applied to the L4 and L2 datasets. BactDating was run in both cases but failed to converge after 1e$^{7}$ or 3e$^{7}$ MCMC iterations for the L4 dataset. The subsampled BEAST2 runs highlighted yellow provided the highest likelihood following path sampling over all strict clock models and were run only on accessions with associated collection dates. These rates were applied to the maximum likelihood phylogenetic tree for temporal estimation of resistance emergence (main text Figure 2, see Methods).

| Lineage | Analysis | Missing dates dropped | | Using est dates | |
|---|---|---|---|---|---|
| | | tmrca | sub rate (subs/site/year) | tmrca | sub rate (subs/site/year) |
| L2 | BactDating 1e7 | 389 (-4201 - 963) | 1.7e-07(1.5e-07 - 2.2e-07) | 469 (-2749 - 894) | 1.8e-07(1.6e-07 - 2.2e-07) |
| | BactDating 3e7 | 367 (-3257 - 987) | 1.8e-07(1.5e-07 - 2.2e-07) | 436 (-3252 - 903) | 1.8e-07(1.58e-07 - 2.28e-07) |
| | BEAST2*, strict coalescent skyline | 453 (-336 - 904) | 7.7e-08 (4.9e-08 - 1.03e-07) | NA | NA |
| | BEAST2*, strict coalescent exponential | 415 (-280 - 861) | 7.4e-08 (4.9e-08 - 9.8e-08) | NA | NA |
| | BEAST2*, strict coalescent constant | 465 (-218 - 877) | 7.7e-08 (5.2e-08 - 1.01e-07) | NA | NA |
| | BEAST2*, relaxed coalescent skyline | 509 (-505 - 1049) | 8.2e-08 (4.8e-08 - 1.12e-07) | NA | NA |
| | BEAST2*, relaxed coalescent exponential | 727 (-52 - 1217) | 7.9e-08 (4.7e-08 - 1.1e-07) | NA | NA |
| | BEAST2*, relaxed coalescent constant | 554 (-311 - 1088) | 8.6e-08 (5.5e-08 - 1.17e-07) | NA | NA |
| L4 | BEAST2*, strict coalescent skyline | 292 (35 - 502) | 7.1e-08 (6.2e-08 - 7.9e-08) | NA | NA |
| | BEAST2*, strict coalescent exponential | 358 (116 - 557) | 7e-08 (6.2e-08 - 7.9e-08) | NA | NA |
| | BEAST2*, strict coalescent constant | 236 (-32 - 450) | 6.9e-08 (6e-08 - 7.7e-08) | NA | NA |
| | BEAST2*, relaxed coalescent skyline | 336 (-20 - 623) | 7.3e-08 (6.2e-08 - 8.3e-08) | NA | NA |
| | BEAST2*, relaxed coalescent exponential | 567 (278 - 794) | 7.2e-08 (6.1e-08 - 8.2e-08) | NA | NA |
| | BEAST2*, relaxed coalescent constant | 224 (-161 - 526) | 6.9e-08 (5.9e-08 - 7.9e-08) | NA | NA |

**Supplementary Table S4:** Classification of previously observed resistance associated with *mmpR5* variants identified in this study.

| Observed resistance | Mutation | Reference |
|---|---|---|
| Resistant | Val1Ala | Bloemberg [1] |
| | Ser2Ile | Xu 2017 [2] |
| | Glu21Asp | Yang 2018 [3] |
| | Gln22Pro | Nimmo 2020 [4] |
| | Ser31Arg | Yang 2018 [3] |
| | Leu32Ser | CRyPTIC, Sonnenkalb 2023 [5] |
| | Thr33Ala | Ismail 2019 [6] |
| | Thr33Ser | Yang 2018 [4] |
| | Ala36Thr | Ismail 2019 [6] |
| | Cys46Arg | Ismail 2019 [6] |
| | Cys46fs | Nimmo 2020 [4] |
| | Pro48Leu | CRyPTIC, Sonnenkalb 2023 [5] |
| | Arg50Trp | Villellas 2017 [7], Sonnenkalb 2023 [5] |
| | Ser52Phe | Villellas 2017 [7] |
| | Ser53Pro | Ismail 2019 [6] |
| | Ala57Glu | Nimmo 2020 [8] |
| | Ala59Val | Villellas 2017 [7] |
| | Ala62Val | Villellas 2017 [7] |
| | Ile67Ser | Ismail 2018 [9] |
| | Ser68Gly | Andries 2014 [10] |
| | Arg72Trp | Ismail 2019 [6], Sonnenkalb 2023 [5] |
| | Arg72Thr | Nimmo 2020 [8] |
| | Leu74Val | CRyPTIC, Sonnenkalb 2023 [5] |
| | Leu74Pro | Ismail 2019 [6] |
| | Phe79Ser | Yang 2018 [3] |
| | Leu83Pro | Ismail 2019 [6] |
| | Ala84Val | Sonnenkalb 2023 [5] |
| | Asp88Arg | Nimmo 2020 [8] |
| | Tyr92Cys | Karmakar 2020 [11] |
| | Arg94Gln | Andries 2014 [10] |
| | Arg96Trp | CRyPTIC, Sonnenkalb 2023 [5] |
| | Arg96Gly | CRyPTIC, Sonnenkalb 2023 [5] |
| | Asn98Asp | Yang 2018 [3] |
| | Asn98fs | Ismail 2019 [6] |
| | Ala102Pro | Ismail 2019 [6] |
| | Ile108Val | Andres 2020 [12] |
| | Ala112Ser | Villellas 2017 [7] |
| | Leu117Arg | Xu 2017 [2] |

| | | |
|---|---|---|
| | Ala118Thr | Nimmo 2020 [8] |
| | Gly121Arg | Nimmo 2020 [8] |
| | Leu122Pro | Nimmo 2020 [8] |
| | Arg123Lys | Yang 2018 [3] |
| | Arg135Gly | Ismail 2019 [6] |
| | Leu136Arg | Nimmo 2020 [8] |
| | Met139Ile | CRyPTIC, Sonnenkalb 2023 [5] |
| | Leu154Pro | Ismail 2018 [9] |
| | Met146Thr | Xu 2017[3], note reported susceptible in Yang 2018 |
| Intermediate | Pro14Leu | CRyPTIC, Sonnenkalb 2023 [5] |
| | Val20Gly | Nimmo 2020 [8] |
| | Leu40Ser | Zimenkov 2017 [13] |
| | Ser53Leu | Pang 2017 [14] |
| | Ile67Val | Yang 2018 [3] |
| | Arg89Trp | Nimmo 2020 [8] |
| | Arg90Cys | Yang 2018 [3] |
| | Phe93Ser | Nimmo 2020 [8] |
| | Asn98Val | Zimenkov 2017 [13] |
| | Arg109Leu | Nimmo 2020 [4] |
| | Glu113Lys | Zimenkov 2017 [13] |
| | Gly121Glu | Zimenkov 2017 [13] |
| | Glu138Gly | Andries 2014 [10] |
| | Met139Thr | Veziris 2017 [15] |
| | Leu142Arg | Zimenkov 2017 [13] |
| Susceptible | Asp5Gly | Martinez 2018 [16] |
| | Met17Leu | Villellas 2017 [7] |
| | Met23Leu | Zimenkov 2017 [13] |
| | Met23Val | Martinez 2018 [16] |
| | Leu40Val | CRyPTIC, Sonnenkalb 2023 [5] |
| | Trp42Arg | Villellas 2017 [7] |
| | Glu55Asp | Martinez 2018 [16] |
| | Gly66Trp | Zimenkov 2017 [13] |
| | Gly78Arg | Zimenkov 2017 [13] |
| | Val85Ala | Zimenkov 2017 [13] |
| | Gly87Arg | Martinez 2018 [16] |
| | Asp88Gly | Yang 2018 [3] |
| | Phe93Leu | Nimmo 2020 [4] |
| | Ala101Thr | Villellas 2017 [7] |
| | Asp116His | Villellas 2017 [7] |
| | Arg135Trp | Zimenkov 2017 [13] |
| | Tyr145Phe | Villellas 2017 [7] |
| | Tyr157Asp | Pang 2017 [14] |

| | | |
|---|---|---|
| | Ser158Arg | Villellas 2017 [7] |
| Hypersusceptible | C-11A | Villellas 2017 [7] |
| Undetermined frameshifts | Asp47fs | Sonnenkalb 2023 [5] |
| | Asp141fs | Unvalidated, Rancoita 2018 [17] indicate susceptible |

**Table S5:** Sequence data identified with *mmpR5* variants predating 2007 with a known phenotype. Dates flagged with an asterisk (*) indicate those dates which have been permuted using the metadata of all samples (see Est. column of Supplementary Table S1).

| **Lineage 2** | | | | |
|---|---|---|---|---|
| **Accession** | **Collection Year** | ***mmpR5* variant** | **Predicted Phenotype** | **Location** |
| **SRR067502** | 2000 | Ala59Val | Resistant | USA |
| **SRR067488** | 2000 | Ala59Val | Resistant | Missing |
| **SRR067486** | 2002 | Ala59Val | Resistant | Missing |
| **SRR067484** | 2003 | Ala59Val | Resistant | Missing |
| **SRR070032** | 2003 | Ala59Val | Resistant | Missing |
| **SRR067534** | 2004 | Ala59Val | Resistant | Missing |
| **SRR6397632** | 2005.5* | C-11A | Hypersusceptible | Canada |
| **SRR1013665** | 2005 | Asn98fs | Resistant | South Korea |
| **SRR6153262** | 2006.5* | C-11A | Hypersusceptible | Canada |
| **SRR067468** | 2006 | Ala59Val | Resistant | Missing |
| **Lineage 4** | | | | |
| **MDRDM260** | 1999 | Ile67fs + mmpL5 R202fs | Hypersusceptible | Missing |
| **ERR028625** | 1999 | Val20Ala | Intermediate | Netherlands |
| **ERR025419** | 1999 | Val20Ala | Intermediate | Netherlands |
| **ERR025417** | 1999 | Val20Ala | Intermediate | Netherlands |
| **ERR025414** | 1999 | Val20Ala | Intermediate | Netherlands |
| **ERR025418** | 1999 | Val20Ala | Intermediate | Netherlands |
| **ERR028626** | 1999 | Val20Ala | Intermediate | Netherlands |
| **LE-006** | 2003 | Ile67fs + mmpL5 R202fs | Hypersusceptible | Peru |
| **HO-109** | 2003 | Ile67fs + mmpL5 R202fs | Hypersusceptible | Peru |
| **LN3756** | 2004 | Ile67fs + mmpL5 R202fs | Unknown | Missing |
| **HO-251** | 2004 | Ile67fs + mmpL5 R202fs | Hypersusceptible | Peru |
| **LN-3755** | 2004 | Ile67fs + mmpL5 R202fs | Hypersusceptible | Peru |
| **ERR2652986** | 2004 | Asp127Val | Unknown | Brazil |
| **PAR-044** | 2005 | Ile67fs + mmpL5 R202fs | Hypersusceptible | Peru |
| **ERR023260** | 2005.73* | Glu28Ala | Unknown | USA |

**Table S6:** Number of estimated emergence (homoplasic) events for major *mmpR5* variants considered.

| Variant | Predicted phenotype | L2 | L4 |
|---|---|---|---|
| Ile67fs | Resistant | 11 | 7 |
| Ile67fs + mmpL5 Arg202fs* | Hypersusceptible | 0 | 1 |
| c-11a | Hypersusceptible | 1 | 0 |
| Asp5Gly | Susceptible | 2 | 1 |
| Met146Thr | Resistant | 1 | 2 |
| Leu40Val | Susceptible | 0 | 1 |
| Glu49fs | Resistant | 2 | 7 |
| Arg90Cys | Intermediate | 1 | 1 |
| Asp47fs | Resistant | 7 | 1 |
| Ala59Val | Resistant | 1 | 1 |
| Asn98Asp | Resistant | 0 | 3 |
| Leu117Arg | Resistant | 2 | 4 |
| Val1Ala | Resistant | 2 | 0 |
| Val20Ala | Intermediate | 1 | 1 |
| Arg156fs | Resistant | 0 | 2 |
| Asp141fs | Resistant | 0 | 1 |
| Gly121Arg | Resistant | 1 | 1 |
| Arg109Leu | Resistant | 0 | 1 |
| Arg96Gly | Resistant | 0 | 4 |

**Table S7 [external excel document]:** Inferred age of nodes and preceding nodes for L2 samples with *mmpR5* nonsynonymous variants. Cells are coloured as per phenotype annotations (see main text Figure 1-3). Presence of mmpL5 variants is noted, as are MICs where available. TBProfiler resistance profiles as either "S" for susceptible, "RR" for rifampicin-resistant and "preXDR" for fluoroquinolone-resistant.

**Table S8 [external excel document]:** Predicted probability of bedaquiline resistance based on the amino acid properties of *mmpR5* variants following a machine learning predictive approach (see Methods and Supplementary Note 1).

**Table S9:** Precision, recall, F1, AUPRC, sensitivity and specificity scores for gradient-boosted tree classifier. Standard deviation was calculated across the 10 outer loops of the nested cross-validation protocol.

| *Metric* | *Score* | *Standard dev.* |
|---|---|---|
| *Precision* | 0.826 | 0.1 |
| *Sensitivity/Recall* | 0.754 | 0.1 |
| *F1* | 0.780 | 0.08 |
| *AUPRC* | 0.787 | 0.1 |
| *Specificity* | 0.633 | 0.3 |

### Supplementary Note 1

#### Methods

A gradient-boosted tree classifier was developed using the XGBoost API (v1.0.2)[20] to predict the phenotypic effect of *mmpR5* variants based on the amino acid properties of known resistant and susceptible variants. Mutations that confer resistance were used as the positive class in this binary classifier. Fifteen features were engineered based on the wild type and mutant residues of each mutation to include the amino-acid residue type, the polarity, charge, hydrophobic or hydrophilic status, molecular weight, and location including the 5'-3' position.

Only variants in *mmpR5* which have a demonstrated association to a bedaquiline-resistant or susceptible phenotype were used, resulting in 55 resistance and 27 susceptibility mutations. Model parameters were optimised to maximise the F1 score and model performance was estimated using a nested, stratified, 10 x 10 cross-validation procedure. AUPRC was used for model evaluation due to class imbalance in the dataset [18]. The trained model was then interpreted using TreeExplainer as part of the shap API (v0.35.0) (64) (**Supplementary Figure S15**) to allow inference of how each feature contributes to each prediction. All scripts used for the analysis are hosted on GitHub (https://github.com/cednotsed/TB-Bedaquiline-Resistance-Modelling.git). Predictions were also made using the Protein Variation Effect Analyzer (PROVEAN) via the online interface [19]. The final predicted probability of resistance and associated PROVEAN scores are provided in **Supplementary Table S8**.

#### Results

To assess properties associated to RAVs which may be useful predictors of the phenotypic effect of these unknown variants we applied a machine learning approach. A gradient-boosted tree classifier (XGBoost API (v1.02) was trained and optimised to determine if the amino acid properties of *mmpR5* mutations associated to known bedaquiline resistance phenotypes can be used to predict the resistance status of mutations with no available phenotypic information.

Fifteen features were engineered based on the wild type and mutant residues of each mutation as follows. Two features represent the amino-acid residue of the wild type and of the mutant. Two features encode whether they are non-polar, polar, positively charged or negatively charged. Two features represent whether the wild type and the mutant residues are hydrophobic or hydrophilic, based on the hydrophobicity scale proposed by Janin[20]. Two features encode the molecular weight of wild type and mutant AA. Two features represent the change in charge or molecular weight from the wild type to mutant, where non-polar and polar residues are assumed to contribute a charge of zero. One feature represents the change in hydrophobicity, where a hydrophobic→hydrophilic residue change is coded as +1 and the reverse as -1. Three features represent mutations in the DNA-binding domain, in the dimerisation domain, and to the residues in contact with 2-stearoylglycerol. The last feature represented the 5'- 3' position of amino acid mutations.

Only variants in *mmpR5* which have a demonstrated association to a bedaquiline-resistant or susceptible phenotype were used, resulting in 55 resistance and 27 susceptibility mutations. In this instance we excluded frameshifts from the analysis as it is challenging to encode their physiochemical characteristics. Model parameters were optimised to maximise the F1 score and model performance was estimated using a nested, stratified, 10 x 10 cross-validation procedure. In this procedure, the model is trained and optimised on a subset of the data and the resultant model is evaluated on another subset of the data not used for training and optimisation. This process is then repeated 10 times and the model performance metrics are averaged to provide an assessment of how well the models perform on unseen data. AUPRC was used for model evaluation due to class imbalance in the dataset[18]. The optimised model provided an area under the precision-recall curve (AUPRC) of 0.79 (**Supplementary Table S8**), suggesting that the physiochemical properties of mutations can be used to successfully differentiate between resistance and susceptibility phenotypes (**Supplementary Table S9**).

The trained model was then interpreted using TreeExplainer as part of the shap API (v0.35.0) (64). Each feature is assigned a SHAP value which represents the change in predicted probability score in each prediction when a feature is included or excluded from the model. Visualisation of the SHAP values in tandem with the feature values (**Supplementary Figure S15**) allows inference of how each feature contributes to each prediction. The features of the models were then interpreted using SHAP values. As a measure of the relative importance and effect sizes for these predictor variables in the model, we calculated the mean absolute SHAP (MAS) values for all predictors, which measures the magnitude of the change in predicted probability of resistance caused by each variable in the model. For example, the variable 'WT hydrophobicity' (i.e., a measure of whether the wild-type amino acid is hydrophobic or not) had a MAS value of 7.8%, indicating that on average, this variable adds or subtract 7.8% to the probability of resistance predicted by the model. Via this approach, we found that mutations to polar and positively-charged residues, or of polar and positively-charged residues, and those to higher molecular weight amino acids are associated with resistance (MAS=2.1%, 4.5% and 2.5%, respectively). Conversely, mutations in the dimerisation or DNA-binding domains, transitions from negatively to positively charged residues, and mutations of hydrophobic residues, or mutations to hydrophobic residues are associated with susceptibility (MAS=4.8%, 2.2%, 2.6%, 7.8%, and 4.3%, respectively; **Supplementary Figure S15**). Overall, we demonstrate the promise of using the physiochemical characteristics of single mutations as input to machine learning models to predict their effects on drug susceptibility. Additionally, we demonstrate the power of model interpretation approaches such as SHAP to derive biological insights from complex machine learning models.

In the absence of phenotypic data, machine learning approaches offer possibilities to predict the resistance status of given variants, and our small-scale analysis suggests the potential of such an approach. Our model uses general physiochemical characteristics of single non-synonymous mutations to infer information about drug susceptibility, which in principle allows us to predict the phenotypic effects of novel mutations that have not been characterised. However, even determining the phenotypic consequences of *mmpR5* variants that have previously been described is challenging as there are often only limited reports correlating MICs to genotypes. Moreover, at least four different methods are used to determine MICs, some of which do not have associated critical concentrations. Even where critical concentrations have been set, there is an overlap in MICs of isolates that are genetically wild type and those that have mutations likely to cause resistance [21]. Further, our machine learning modelling approach based on amino acid properties alone does not consider epistatic interactions that arise from the genetic background of isolates such as the mmpL5 frameshift mutations we observed. Future work aiming to include more genetic information in machine learning models, including frameshift mutations or different combinations of point mutations, may help better understand the correlations between genotype, protein function and phenotype, and may ultimately lead to better resistance predictions.

All scripts used for the analysis are hosted on GitHub (https://github.com/cednotsed/TB-Bedaquiline-Resistance-Modelling.git). Predictions were also made using the Protein Variation Effect Analyzer (PROVEAN) via the online interface[22]. The final predicted probability of resistance and associated PROVEAN scores are provided in **Supplementary Table S8**.
